# Platelets exploit fibrillar adhesions to assemble fibronectin matrix revealing new force-regulated thrombus remodeling mechanisms

**DOI:** 10.1101/2020.04.20.050708

**Authors:** Sebastian Lickert, Kateryna Selcuk, Martin Kenny, Johanna L. Mehl, Susanna M. Früh, Melanie A. Burkhardt, Jan-Dirk Studt, Ingmar Schoen, Viola Vogel

**Affiliations:** Laboratory of Applied Mechanobiology, Department of Health Sciences and Technology, ETH Zurich, Vladimir-Prelog-Weg 4, 8093 Zurich, Switzerland; Irish Centre for Vascular Biology, School of Pharmacy and Biomolecular Sciences, Royal College of Surgeons in Ireland, 123 St Stephen’s Green, Dublin 2, Ireland; Hahn-Schickard, Georges-Koehler-Allee 103, 79110 Freiburg, Germany & Laboratory for MEMS Applications, IMTEK Department of Microsystems Engineering, University of Freiburg, Georges-Koehler-Allee 103, 79110 Freiburg, Germany; Division of Hematology, University Hospital Zurich, Rämistrasse 100, 8091 Zurich, Switzerland

## Abstract

Upon vascular injury, platelets are crucial for thrombus formation and contraction, but do they directly initiate early tissue repair processes? Using 3D super-resolution microscopy, micropost traction force microscopy, and specific integrin or myosin IIa inhibitors, we discovered here that platelets form fibrillar adhesions. They assemble fibronectin nanofibrils using αIIbβ3 (CD41/CD61, GPIIb-IIIa) rather than α5β1 integrins, in contrast to fibroblasts. Highly contractile platelets in contact with thrombus proteins (fibronectin, fibrin) pull fibronectin fibrils along their apical membrane, whereas platelets on basement membrane proteins (collagen IV, laminin) are less contractile generating less stretched planar meshworks beneath themselves. As probed by vinculin-decorated talin unfolding, platelets on fibronectin generate similar traction forces in apical fibrillar adhesions as fibroblasts do. These are novel mechanobiology mechanisms by which platelets spearhead the fibrillogenesis of the first *de novo* ECM, including its 2D versus 3D network architectures depending on their ECM environment, and thereby pave the way for cell infiltration.

## INTRODUCTION

When blood vessels get injured, the massive recruitment and activation of platelets culminates in thrombus formation to stem bleeding, whereas the subsequent infiltration of the clot by immune and stromal cells is essential to ultimately restore tissue integrity (Rodrigues et al. 2019). Platelet adhesion receptors initiate interactions with extracellular matrix (ECM) proteins that become exposed by endothelial barrier disruption, while platelet signaling receptors boost platelet aggregation and granule secretion. Both signal types guide the adoption of several different platelet phenotypes at different times and locations in the thrombus (van der Meijden and Heemskerk 2019; Brass, Diamond, and Stalker 2016). The ‘pro-hemostatic’ secretion of von Willebrand Factor (vWF), fibrinogen (Fg), and fibronectin (Fn) occurs predominantly at injury site boundaries and in the extravascular portion of the plug (Tomaiuolo et al. 2019). Fibrin(ogen)-binding platelets adopt an aggregating phenotype, while the ‘procoagulant’ phosphatidylserine (PS) exposing platelets aid the activation of clotting factors (Schoenwaelder et al. 2009) that in turn catalyze the conversion of fibrinogen to fibrin (Fb) and fibrin crosslinking. This enables the build-up of a complex and anisotropic clot architecture (Brass, Diamond, and Stalker 2016). While the molecular regulation of individual components of the hemostatic system has been studied in detail, how to integrate the many fine-tuned platelet-responses at the systems level to ultimately initiate tissue healing is poorly understood, especially with regards to the role played by the rapidly changing extracellular microenvironment (Brass, Diamond, and Stalker 2016; Tomaiuolo et al. 2019).

Studying the response of platelets to free-floating ECM proteins in solution, where these lack their material properties as interconnected networks, has long hampered the elucidation of platelet mechanobiology (Hansen et al. 2018). Platelets contract rapidly when they get in contact with immobilized fibrinogen and reach maximal forces within 15 minutes (Hansen et al. 2018). Actomyosin-based contractility mediates the probing and sensing of substrate stiffness, which modulates platelet activation (Hansen et al. 2018; Zhang et al. 2018) and drives the bundling and alignment of fibrin fibers (Ono et al. 2008) and thus, clot compaction. Contractile platelets then expel the non-contractile, non-adhesive procoagulant platelets to the thrombus surface (Nechipurenko et al. 2019). Graded platelet contractility is thus key to orchestrate fibrin clot remodeling and the local packing density of platelets within the developing thrombus (Brass, Diamond, and Stalker 2016). The inverse question, whether and if so, how platelet contractility is regulated by the microenvironment inside the thrombus or at injury site boundaries in contact with basement membrane debris, is not well understood because a direct comparison of platelet traction forces on different ECM proteins is missing. This question is clinically most relevant because defects in platelet constituents that mediate contractility (adhesion receptors, actin cytoskeleton, myosin IIa) result in compromised thrombus formation, thrombus instability, and bleeding (Ono et al. 2008; Ting et al. 2019; Myers et al. 2017; Nurden and Nurden 2008).

After its crucial role to stop bleeding is fulfilled, the fibrin clot has to be gradually replaced with newly assembled ECM fibrils (Burkhardt et al. 2016). It is neither known how this is accomplished without compromising its mechanical stability, nor how the composition and architecture of the thrombus regulate its infiltration by surrounding or homed-in cells. Even the question whether platelets not only secrete ECM proteins, but also contribute to the first steps of tissue repair through assembly of the first provisional Fn matrix is controversial. While plasma fibronectin (pFn) contributes to hemostasis and affects thrombus structure (Wang et al. 2014), and platelet-deposited Fn is deoxycholate-insoluble (Olorundare et al. 2001; J. Cho and Mosher 2006b, 2006a) which suggests that it is in its fibrillar form, the spatial resolution of confocal microscopy falls short to confirm the dimensions of Fn fibrils, while few electron microscopy data are available that suggest that fibrils might be as thin as 20 nm (Olorundare et al. 2001). At first glance surprising is also why many different ECM protein coatings support Fn deposition by platelets, including Fn and fibrin (Olorundare et al. 2001), laminin (J. Cho and Mosher 2006b), or collagen type I (Jaehyung Cho and Mosher 2006), whereas others such as vitronectin or fibrinogen (Jaehyung Cho et al. 2005) or von Willebrand factor (vWF) (Jaehyung Cho and Mosher 2006) do not. Specific inhibitors against integrins αIIbβ3 or α5β1 reduced Fn deposition on Fn or fibrin substrates to different degrees (Olorundare et al. 2001), but it remained again unclear whether this reduction was due to reduced platelet-substrate adhesion or to reduced Fn fibrillogenesis. Even for fibroblasts, the molecular mechanism how their mechanobiology drives the formation of fibrillar adhesions (Geiger et al. 2001) is still debated and refined (Lu et al. 2020), and nothing is known about the molecular machinery that drives Fn fibrillogenesis in platelets.

Here we employed direct stochastic optical reconstruction microscopy (dSTORM) to resolve the 3D ultrastructure of platelet-matrix adhesion sites, and micropost arrays to quantify how the adhesion site ultrastructure correlates with platelet traction forces exerted on different ECM protein coatings. We discover that platelets can form fibrillar adhesions too, determine conditions under which they form and the integrins involved. We show that platelets assemble the first provisional Fn matrix with fibers that form an in-plane network when platelets are exposed to proteins, while they assemble a 3D Fn network when exposed to blood clot proteins. These findings point to so far unrecognized mechanisms how sensing and responding to their microenvironment is controlled by platelet contractility and we suggest that this contributes to the emergence of the structural heterogeneity of a thrombus. Our findings also challenge the common notion that the infiltration of fibroblasts or mesenchymal stem cells (Rodrigues et al. 2019) is directed by the fibrin scaffold and that these infiltrating cells are the first to assemble a *de novo* Fn ECM (Singh, Carraher, and Schwarzbauer 2010; Barker and Engler 2017).

## RESULTS

### Substrate adherent platelets assemble fibronectin nanofibrils

Since confocal microscopy falls short to characterize platelet-assembled Fn fibrils (Olorundare et al. 2001; J. Cho and Mosher 2006b, 2006a), we used super-resolution (3D dSTORM) microscopy to analyze washed human platelets on Fn-coated glass and supplemented the medium with fluorescently labeled human pFn that is harvested by cells for fibrillogenesis (G. Baneyx, Baugh, and Vogel 2002) (Figure 1a). Fn fibrils were formed at the platelet periphery mainly at two ends (Figure 1b and Supplementary Figure S1). Fibrils shorter than one micron were difficult to distinguish unambiguously from deposited aggregates and were thus not investigated further. A statistical analysis of longer fibrils (Supplementary Figure S2) revealed that platelets from different normal healthy donors (Supplementary Figure S3) assembled fibrils with consistent characteristic dimensions. Fn fibrils were few microns long (L=1.7±0.43 µm; Figure 1e) and between 20…200 nm thick (D=85±34 nm; Figure 1f). Fibrils were anchored outside of the platelet at the coverslip surface and stretched straight upwards over the platelet lamellipodium (Figure 1b-d) with a mean start-to-end height difference of H=243±123 nm and an inclination of α=8.9±4.9° (Figure 1g,h). Using the same experimental approach, we see that fibroblasts on Fn (Figure 1i,j) produced longer and thicker Fn fibers due to profilament bundling (Früh et al. 2015) (Figure 1k) but showed the same start-to-end height (Figure 1l).

**Figure 1.**
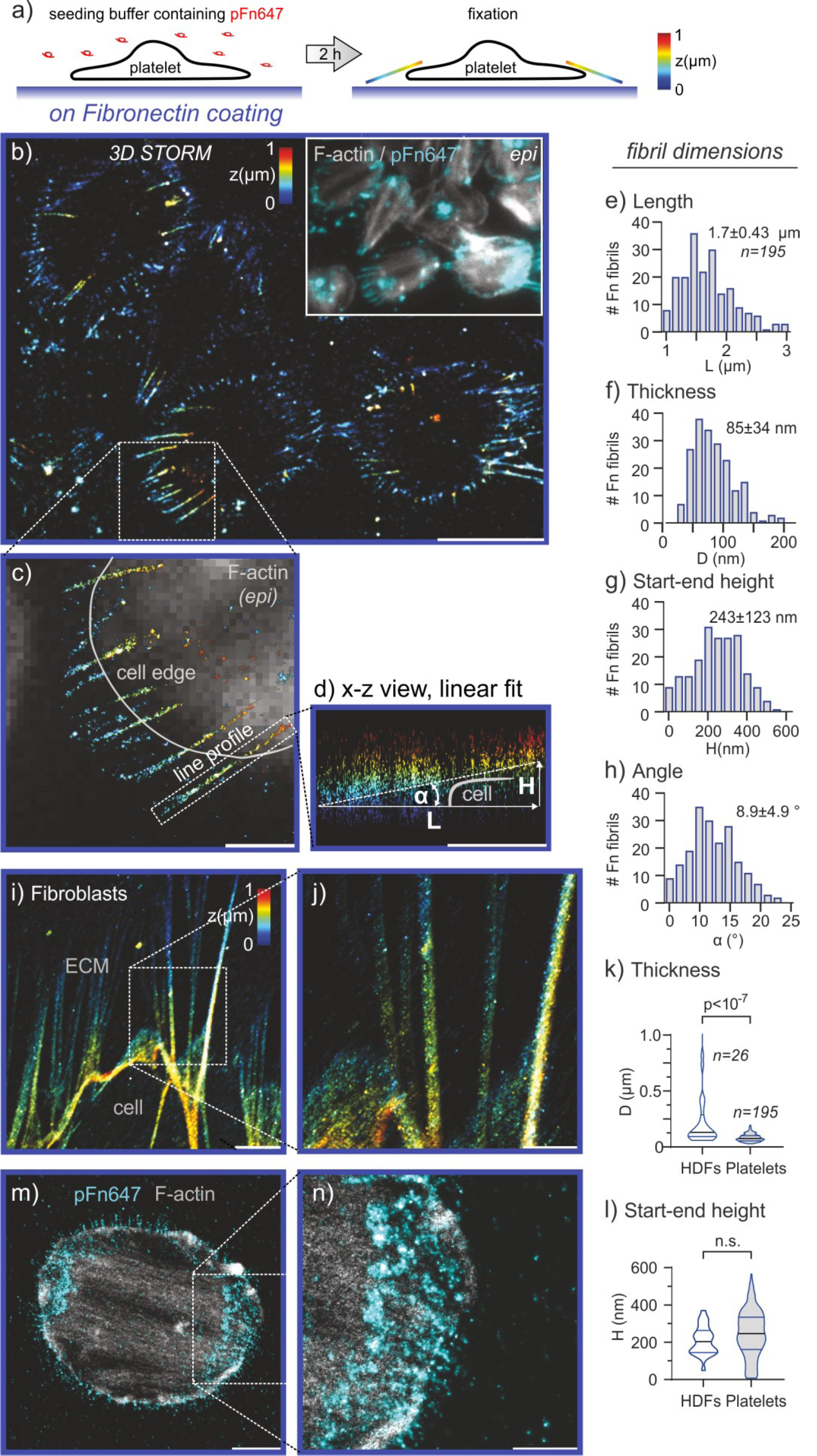
Fibronectin extracellular matrix assembled by human platelets versus by fibroblasts seeded on fibronectin-coated coverslips. **(a)** Sketch of the experimental procedure. Platelets are seeded on fibronectin (Fn, blue)-coated glass coverslips and the medium was supplemented with 90 µg/ml pFn and 10 µg/ml pFn647 for 2 hours at 37°C. The platelets incorporated pFn647 during this time period into their newly assembled Fn fibers. The samples were then fixed for super-resolution imaging. **(b)** pFn647 imaged by 3D dSTORM. The z-position is color-coded from blue (basal) to red (apical). Scale bar 5 µm. Inset: epifluorescence (epi) image of F-actin (grey) and pFn (cyan) of the same cells. **(c)** Magnification of the boxed region of Fn fibrils at the cell edge in (b). Scale bar 1 µm. **(d)** x-z side view of a single Fn fibril (see boxed region in c). The dashed line depicts the linear fit of the localizations of this fibril and its resulting length (L), the start-to-end height (H), and the angle (α) towards the coverslip. Scale bar 1 µm. **(e-h)** Dimensions of 195 Fn fibrils extracted from 3D dSTORM images. Data were pooled from five different healthy donors (31-35 years). Only fibrils longer than 1 µm were included in the analysis. **(i)** 3D dSTORM image of Fn-fibrils assembled by human foreskin fibroblasts (HFFs). Scale bar 5 µm. **(j)** Magnification of the boxed region of Fn fibrils at the cell edge in (i). Scale bar 2 µm. Comparison of **(k)** thickness D and **(l)** start-end height H between platelets and fibroblasts. Data distributions are depicted as violin plots showing the median (thick black line) and interquartile ranges (thin colored lines). Data were compared with an unpaired two-tailed Mann-Whitney test. Adjusted p-values below 0.001 were accepted as highly significant. n.s.: not significant. **(m)** Representative dual color (2C) dSTORM image of a single platelet spread on Fn with supplemented pFn (cyan) during seeding and stained in addition for F-actin (gray). Scale bar 2 µm. **(n)** Magnification of the boxed region of Fn fibrils at the cell edge in (m). Scale bar 1 µm.

### Platelet Fibronectin fibrils align with tensed and polarized actin fiber bundles

As Fn fibrillogenesis is initiated by tensile forces that partially expose cryptic Fn-Fn assembly sites (Hocking, Smith, and McKeown-Longo 1996; Gretchen Baneyx and Vogel 1999) and an impaired cytoskeleton reduces Fn deposition by platelets (J. Cho and Mosher 2006b) and by other cells (G. Baneyx, Baugh, and Vogel 2002), we next analyzed the spatial organization of the actin cytoskeleton using dual color (2C) dSTORM. Confirming previous observations (Lickert et al. 2018), we found that the majority of platelets on Fn coatings had formed a highly aligned ‘bipolar’ filamentous (F-)actin network with pronounced bundles traversing the cell (Figure 1m and Supplementary Figure S1). Longer Fn fibrils emanated parallel to these actin cables at adhesion sites situated at both ends and extended beyond the platelet edge (Figure 1n).

The linear co-alignment of Fn fibrils with F-actin was similar to fibrillar adhesions in nucleated cells (Zamir et al. 2000). In contrast to focal adhesions which only form on the basal side of fibroblasts in 2D cell culture, fibrillar adhesions in fibroblasts are elongated, fibronectin-associated adhesion structures on either the apical or basal side (Katz et al. 2000). Depending on cell type and time, their composition can differ. By forming fibrillar adhesions, α5β1 integrins are the main drivers of Fn assembly in fibroblasts, yet αvβ3 can partially compensate in the absence of α5β1 (Leiss et al. 2008), and αIIbβ3 integrins overexpressed in CHO cells also mediate Fn assembly (Wu 1997). Emerging fibrillar adhesions in fibroblasts contain talin, which binds to the NPxY motif of β–integrins as well as to actin, while over time, talin is being replaced to variable extents by tensin (Katz et al. 2000) controlled by phosphorylation of the NPxY motif (Legate and Fassler 2009). Very recent evidence suggests the existence of an alternative class of fibrillar adhesions in nucleated cells formed on basement membrane proteins through a mechanism in which an early focal adhesion becomes a sliding fibrillar adhesion that deposits Fn fibrils directly onto the basement membrane (Lu et al. 2020). Here we now show striking morphological similarities between the adhesions fibroblasts and platelets exploit to pull fibronectin fibers, whereby their morphology gave them their name “fibrillar adhesions”. In contrast to fibroblasts, we see little tensin in the fibrillar adhesions of platelets that were fixed 2 hours after seeding (Supplementary Fig. S4). Future research has to elucidate the time-evolving molecular composition of fibrillar adhesions in platelets.

### Fibronectin fibrils are mostly pulled along the apical membrane of platelets spread on fibronectin, while pulled along their basal side on laminin coatings

Since platelets sense different ECM proteins after vascular injury, we asked whether the spatial architecture of *de novo* assembled Fn fibrils depends on the ECM proteins to which the platelets adhere. Fn fibrils were radially oriented on laminin (Ln), in agreement with previously reported Fn deposition patterns (J. Cho and Mosher 2006b), while the actin cytoskeleton showed a rather ring-like arrangement (Lickert et al. 2018) (Supplementary Figure S5). In contrast to platelets on Fn that pulled Fn fibrils along their apical side (Figures 2a-c), platelets on laminin assembled Fn fibrils at their basal side, i.e. between the platelets and the coverslip (Figure 2d-f). Fibrils on laminin were of similar length (L=1.8±0.56 µm) as on Fn and slightly thicker (D=105±39 nm, p<10^−5^), while the start-to-end height (H=90±77 nm, p<10^−15^) and tilt (α=3±2.84°, p<10^−15^) were significantly decreased (Figure 2g-i and Supplementary Figure S5). No statistically significant differences were found for the tested laminin isoforms 111 or 521 (Supplementary Figure S5). We also observed several curvy Fn fibrils on laminin (Figure 2e, arrow) which might indicate that they are less tensed. To validate that most Fn fibrils were attached to the basal platelet membrane, we measured the height of the lamellipodium by imaging the peripheral actin rim at the cell edge and obtained ∼200 nm (Supplementary Figure S6, yellow background in Figure 2i). The fraction of Fn fibrils with H >200 nm was 71% on Fn coatings, in contrast to 9% on laminin coatings. To differentiate between these two distinct Fn fibril architectures, we will refer to them as 3D (on Fn) and 2D (on laminin).

**Figure 2.**
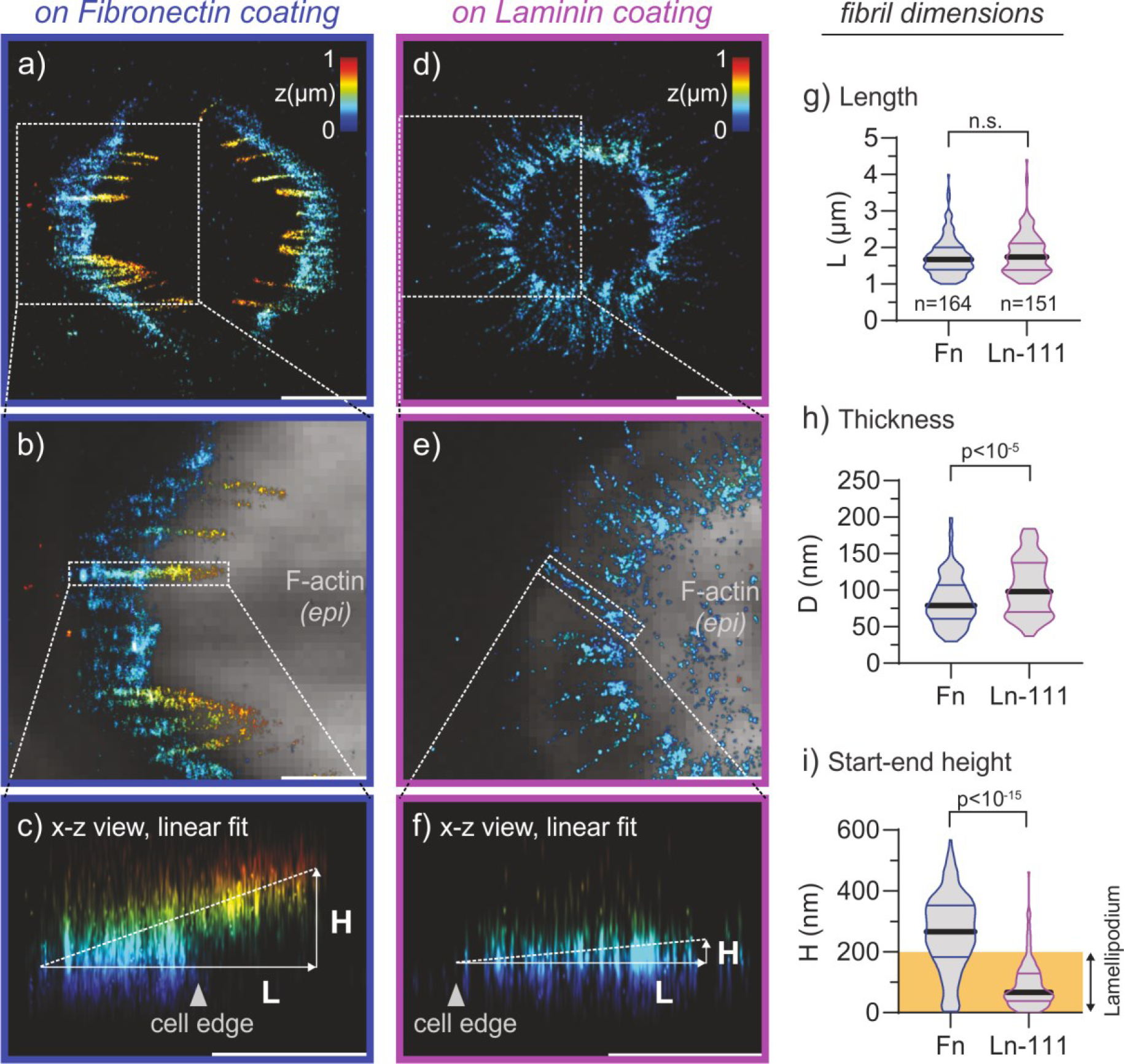
Fibronectin extracellular matrix assembled by human platelets seeded on laminin coatings. See also Supplementary Movie S1. **(a)** dSTORM image of Fn fibrils assembled in the presence of supplemented pFn647 by a representative platelet spread on Fn. Scale bar 2 µm. **(b)** Magnification of the boxed region of Fn fibrils at the cell edge in (a). The dSTORM image is overlaid with the epi image of F-actin (gray). Scale bar 1 µm. Side view of a single Fn fibril (boxed region in b) and linear fit (dashed line). Scale bar 1 µm. **(d)** Fn fibril assembly by a representative platelet spread on a laminin 111 (Ln-111)-coated coverslip. **(e)** Magnification of the boxed region of Fn fibrils at the cell edge in (d). Note the non-straight appearance of Fn fibrils often observed on Ln-111. **(f)** Side view of a single Fn fibril (boxed region in e). Scale bar 1 µm. **(g-i)** Comparison of Fn fibril dimension on Fn-coatings (blue) and Ln-111 (magenta). The orange box in (i) denotes the height of the platelet lamellipodium (see Supplementary Figure S6). Data were pooled from four healthy human donors (27-35 years); for donor-to-donor variation on Fn coatings, see Supplementary Figure S3. Data were compared with an unpaired two-tailed Mann-Whitney test.

### αIIbβ3 integrins are the sole drivers of fibronectin fibrillogenesis in platelets, not α5β1 integrins

Since α5β1 integrins drive Fn fibrillogenesis in mesenchymal cells, yet the platelet integrin αIIbβ3 recognizes Fn’s synergy site (Chada, Mather, and Nollert 2006) too, we next asked which integrin was responsible for Fn fibrillogenesis. Both integrins are expressed at very different levels, with α5β1 counts being 2-3% compared to αIIbβ3 (Zeiler, Moser, and Mann 2014). Since inhibition of αIIbβ3 or α5β1 would also interfere with the anchorage to Fn coatings, we conducted the experiments on laminin where platelets adhere independently via α6β1 (Figure 3a). Blocking αIIbβ3 integrins by the non-priming inhibitor RUC-2 (Zhu et al. 2012) completely abolished Fn fibril formation (Figure 3b). In contrast, blocking α5β1 by the antibody JBS5 (Li et al. 2003) (Figure 3c) showed no differences to the control with regards to the fraction of platelets which assembled Fn fibrils or to fibril length (Figure 3d,e). Note that neither inhibitor affected platelet spreading (Supplementary Figure S7). In contrast to mesenchymal cells, α5β1 is thus dispensable for Fn assembly by platelets and cannot rescue fibrillogenesis in the absence of αIIbβ3 integrins.

**Figure 3.**
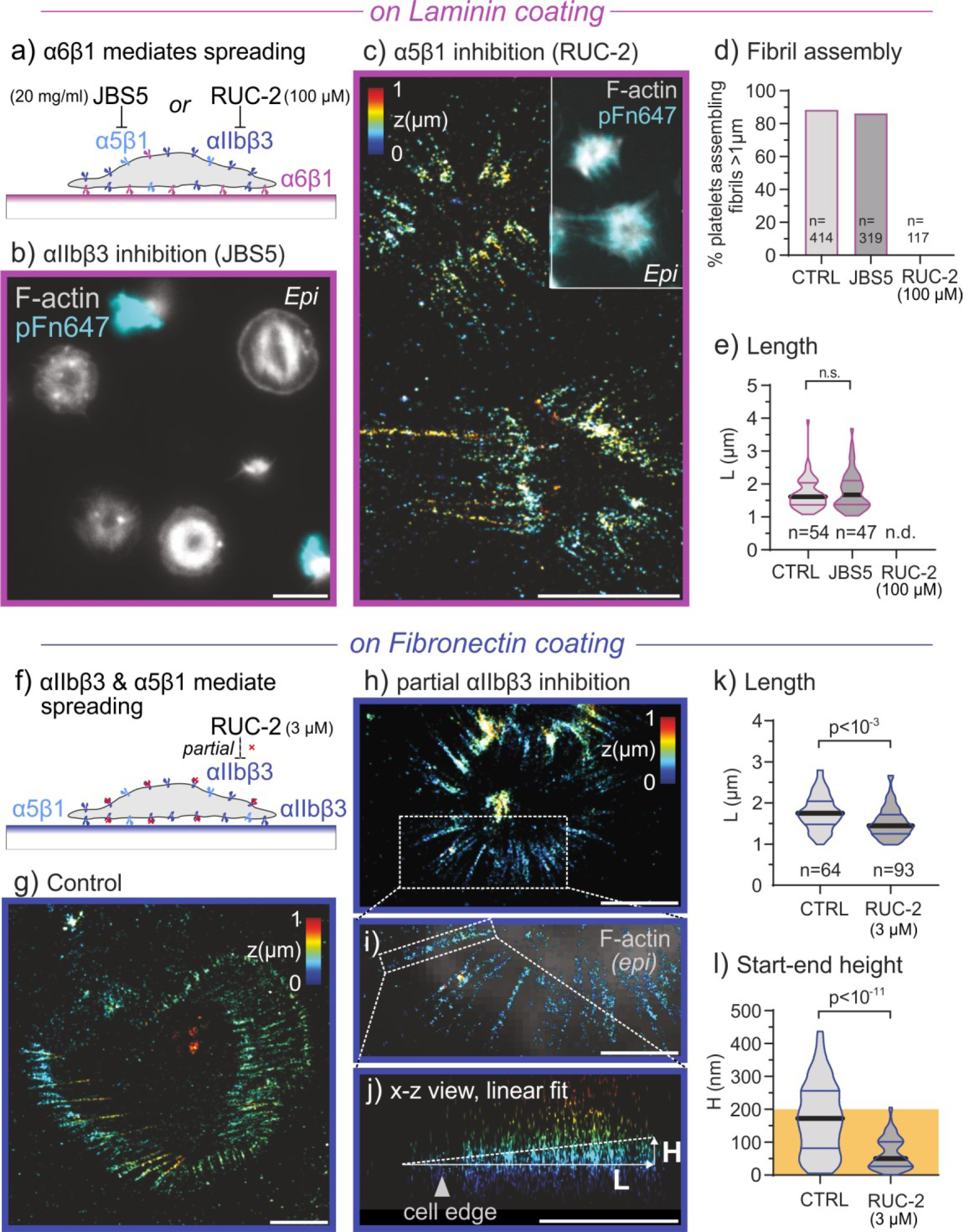
Inhibition of the integrin α5β1 and αIIbβ3 during platelet seeding. **(a)** Sketch of the experimental approach. A high concentration of either the blocking antibody JBS5 against α5β1 (20 mg/L) or of the non-priming inhibitor RUC-2 against αIIbβ3 (100 µM) were supplemented during platelet seeding on Ln-111-coated glass coverslips to block Fn binding through α5β1 and αIIbβ3, respectively. Laminin allows for platelet spreading through integrin α6β1 independently of the blocked integrins. **(b)** Epifluorescence image of F-actin (grey) and pFn647- enriched Fn fibrils (cyan) of platelets inhibited with RUC-2 (100 µM). Scale bar 5 µm. **(c)** 3D dSTORM image of platelets inhibited with JBS5 (20 mg/L). White box: epi image of F-actin (grey) and pFn (cyan) of the same cells. Scale bar 5 µm. **(d)** Comparison of the fraction of platelets that produced micron-long fibrils treated with JBS5 and RUC-2, respectively. Epifluorescence images with the indicated number of cells were analyzed. **(e)** Comparison of Fn fibril length for platelets treated with JBS5 and RUC-2, respectively. Data were obtained from two healthy donors (31 and 33 years). Note that no fibrils were detected for RUC-2 (n.d.: not detectable). **(f)** Sketch of the experimental approach. A sub-saturating concentration (3 µM) of the non-priming αIIbβ3 inhibitor RUC-2 was supplemented during platelet seeding to partially block Fn binding of αIIbβ3 on Fn-coated glass coverslips. **(g)** 3D dSTORM image of platelet assembled Fn fibrils in the presence of pFn647 under control conditions. Scale bar 2 µm. **(h)** 3D dSTORM image of Fn fibrils assembled in the presence of 3 µM RUC-2. Scale bar 2 µm. **(i)** Magnification of the boxed region in (h). The 3D dSTORM image is overlaid with the epi image of F-actin (gray). Scale bar 1 µm. **(j)** Side view of a single Fn fibril (boxed region in i). **(k-l)** Comparison of fibril dimensions. Data were obtained from one male healthy donor (31 years) treated with RUC-2. Data were compared with an unpaired two-tailed Mann-Whitney test.

Interestingly, partial inhibition of αIIbβ3 by low dose RUC-2 still allowed normal platelet spreading on Fn coatings (Supplementary Figure S7) yet had a major impact on Fn fibrils which were now pulled along the basal and not along the apical membrane (Figure 3h-j), very similar to laminin coatings. Thus, reducing the density of functional αIIbβ3 integrins on the platelet surface switched the dimensionality of the pulled fibrils from 3D to 2D.

### Platelet contractility is significantly higher on fibronectin than on laminin coatings

We next asked how the observed phenomena relate to platelet contractility. We thus measured traction forces generated by single platelets using micropost arrays that were coated with Fn or laminin by microcontact printing and then passivated to restrict adhesion of platelets to the functionalized post tops (Figure 4a). Platelets spread similarly on either coating and bent posts towards the center of the cell (Figure 4b-d). A quantification of post deflections yielded higher tractions on Fn compared to laminin, with 2.5±1.0 nN versus 1.9±1.1 nN mean force per post, or 18.7±9.4 nN versus 15.7±9.4 nN per cell, respectively (Figure 4e). The total force per cell as distributed over many posts on Fn was comparable to individual platelets pulling at two anchoring points, i.e. at a suspended AFM tip and a fibrinogen-coated substrate (Lam et al. 2011) or two nearby fibrinogen dots on a compliant polyacrylamide hydrogel (Myers et al. 2017).

### Elevated platelet traction forces are required to induce 3D fibronectin fibril assembly

To test whether a modulation of contractility was sufficient to explain the differences in Fn fibrillogenesis, we used blebbistatin (BBT) which inhibits human myosin IIa with an IC_50_ of 5.1 µM (Limouze et al. 2004). Blocking contractility with high dose BBT (20 µM) led to lessened F-actin bundling and nearly completely abolished the formation of Fn fibrils (Figure 4f). BBT concentrations between 0.3 µM and 10 µM dose-dependently reduced the fraction of fibril-forming platelets from 80% to 20% (Figure 4g), but hardly affected fibril diameter and length (Supplementary Figure S8). Interestingly, fibrils were still anchored at the apical platelet membrane, irrespective of BBT dose (Figure 4h). 3 µM and 10 µM BBT significantly reduced mean platelet traction forces but still showed a minor fraction of platelets with elevated contractility (Figure 4i). Hypothesizing that these contractile platelets were responsible for the residual Fn fibril assembly, we used the measured fraction of fibril-forming platelets at 0, 3, and 10 µM BBT (Figure 4g) and derived a corresponding minimum threshold force from the measured distribution of traction forces, with approximately 0.8 nN, 0.9 nN, and 1.3 nN, respectively (Figure 4i). The good agreement suggests that a minimum traction force of ∼1 nN per cell-substrate adhesion sites is necessary to induce Fn fibrillogenesis in 3D, and that myosin inhibition affects Fn fibrillogenesis in single platelets in an all-or-none fashion, in contrast to the gradual effect of adhesion receptor inhibition (cf. Figure 3h-l).

### The mechanomolecular strain in fibrillar adhesions is the same in fibroblasts and platelets on fibronectin coatings

To determine whether the different forces in cell-substrate adhesions also result in different forces in fibrillar adhesions, we used 2C dSTORM to characterize the molecular build-up of the latter. In fibroblasts, fibrillar adhesions are formed as Fn-bound integrins are coupled via talin to actin fibers which are pulled via myosin II towards the cell center inducing fibrillogenesis along the pulling direction (Geiger et al. 2001). Thereby, talin gets stretched and partially unfolds, opening up binding sites for vinculin which re-enforces these molecular linkages during adhesion maturation (Atherton et al. 2016; Huang et al. 2017). Talin unfolding is accompanied by an increasing spatial offset between integrins and vinculin (Case et al. 2015; Liu et al. 2015). In platelets spread on Fn, vinculin was strongly localized to peripheral adhesion sites and formed string-like patterns that were co-aligned with F-actin bundles (Lickert et al. 2018). Fn fibrils overlapped with these elongated vinculin patterns and were tightly registered (Figure 5a,b), similar to fibrillar adhesions in fibroblasts (Zamir et al. 2000). Analysis of the localization density along linear vinculin-Fn patterns (Supplementary Figure S9), where vinculin decorates stretched talin, revealed a mean spatial offset of 381±195 nm between the terminal ends of Fn fibrils and vinculin patches (n=60; Figure 5g,h). A similar mean offset of 375±278 nm is seen for fibroblasts on Fn coatings (Figure 5e,f,h). This suggests that the mechanical strain of the integrin-talin-vinculin-actin connections that are part of force-bearing apical fibrillar adhesion sites in platelets and apical fibrillar adhesion sites in fibroblasts are similar. On laminin coatings, the offset between vinculin and Fn was only 157±90 nm (n=64; Figure 5c,d,h) and thus significantly smaller. The observed offset range is consistent with measured force-induced talin rod extensions of 60…350 nm (Margadant et al. 2011) and the measured shallow orientation ∼15° of talin within adhesions (Liu et al. 2015).

**Figure 4.**
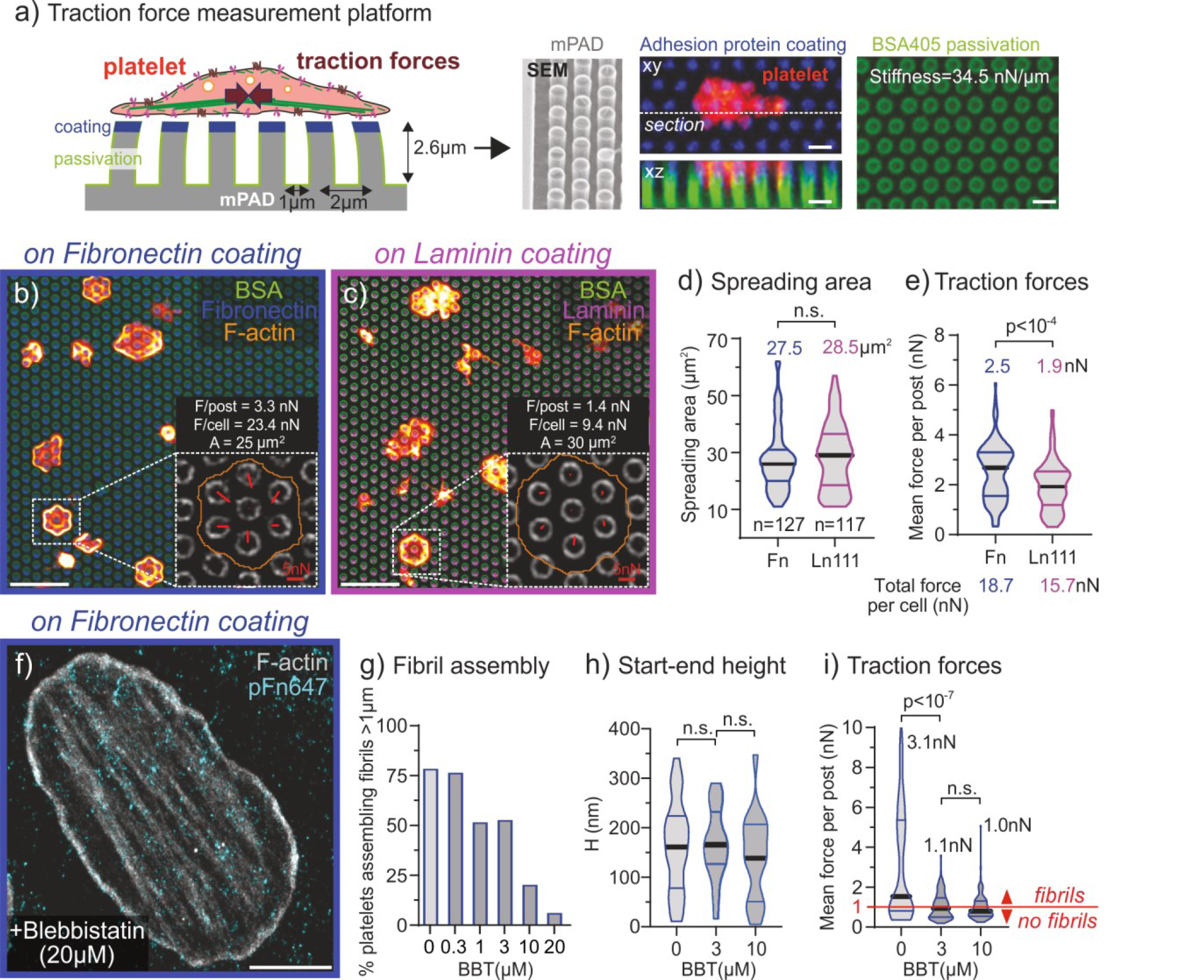
Platelet contractility measured with micropost array detectors (mPADs). **(a)** Traction forces of single platelets are measured with an optimized micropost array. The tops of the posts are stamped with the respective proteins (blue) and the side walls are passivated (green). Post deflections are determined from the centroids of posts in confocal slices using fluorescently labeled BSA (green). **(b**,**c)** Platelets are seeded for one hour on the posts arrays on (b) Fn or (c) Ln-111 and stained for F-actin (red-yellow). Scale bar 5 µm. Insets: force distribution (red arrows) and measured values of a representative platelet (white boxed region). Scale bars 5 nN. **(d**,**e)** Comparisons between Fn and Ln-111 in terms of (d) spreading area on posts or of (e) the mean force per post and total force per platelet. Platelets were obtained from a healthy male donor (33 years) and data were compared with an unpaired two-tailed Mann-Whitney test. **(f)** Representative 2C dSTORM image of a platelet seeded on Fn in the presence of the myosin inhibitor blebbistatin (BBT, 20 µM). As Fn fibril assembly is mostly stalled, pFn647 is seen as cyan dots. Scale bar 2 µm. **(g-i)** Dose dependent effects of BBT on (g) the fraction of platelets that produced micron-long fibrils, (h) fibril start-end height H and (i) mean force per post. The red line in (i) indicates the contractile force level that platelets have to overcome to form Fn fibrils as derived from these experiments. Platelets were obtained from two healthy male donors (31 and 33 years) and data were compared with a non-parametric Kruskal-Wallis rank test with post-hoc Dunn test.

**Figure 5.**
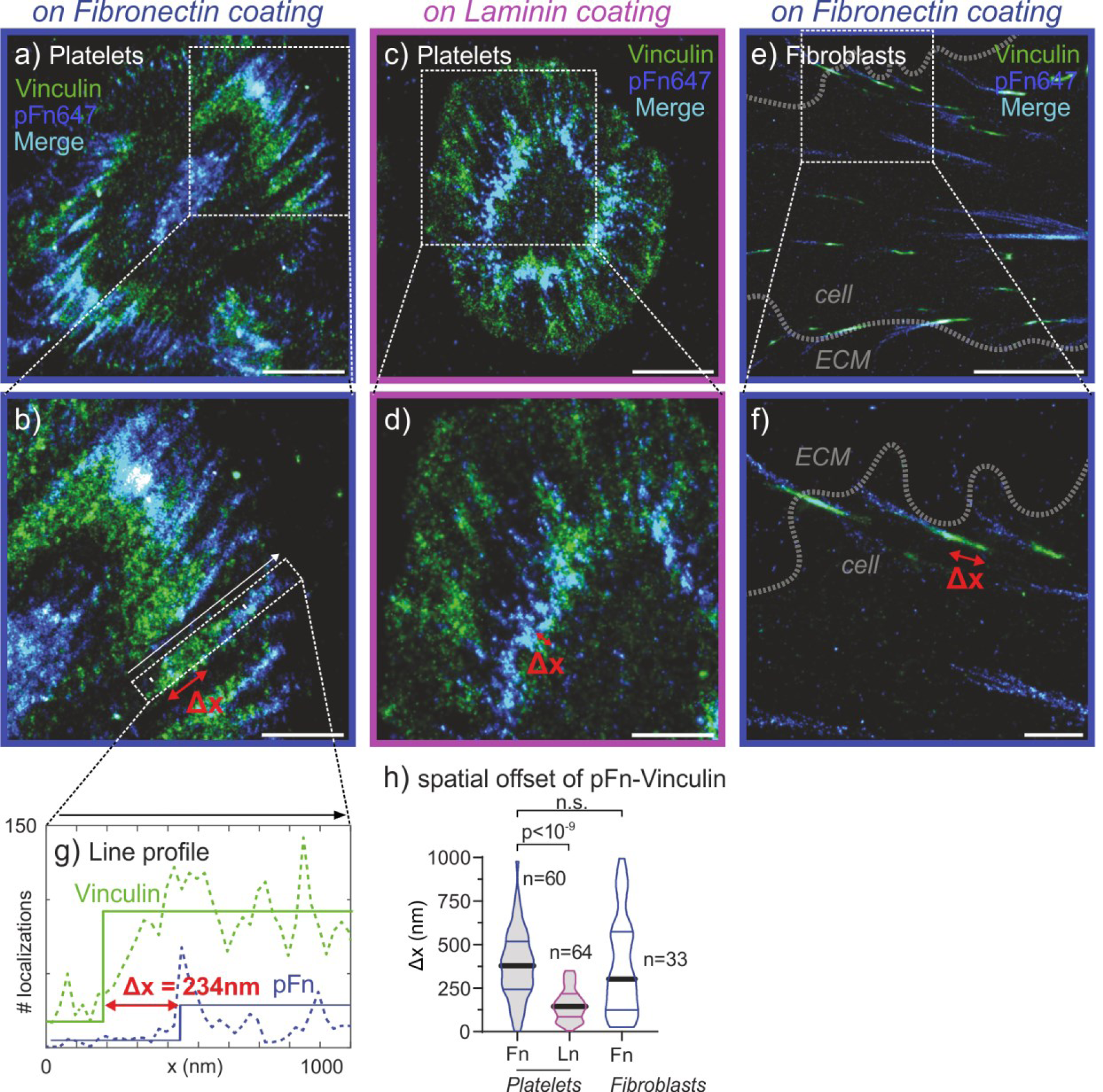
Dual-color dSTORM of fibrillar adhesions in platelets and fibroblasts. **(a)** A representative platelet spread on a Fn coating with medium supplemented pFn (blue) that gets incorporated into the platelet-assembled Fn fibers. The samples were then fixed and stained in addition for vinculin (green). Scale bar 2 µm. **(b)** Magnification of the boxed region of Fn fibrils at the cell edge in (a). Scale bar 1 µm. Red arrow denotes the spatial offset of the vinculin stain with respect to the Fn fibril. **(c**,**d)** A representative platelet spread on Ln-111. Representation as in (a,b). **(e**,**f)** A representative fibroblast seeded on Fn. Scale bars: (e) 5 µm, (f) 1 µm. **(g)** Exemplary line profiles (dashed) of the localization density along the line in (b) for the vinculin stain (green) and the pFn stain (blue). Each profile was fitted with a step function to obtain the spatial offset between stains. **(h)** Comparison of the spatial offset between Fn fibrils and vinculin for platelets on Fn and Ln-111 and for fibroblasts on Fn. Platelets on Fn and Ln were obtained from two same donors, a healthy male and a female donor (30 years and 27 years). Data were compared with a non-parametric Kruskal-Wallis rank test with post Dunn test to make (multiple) comparisons.

### The dimensionality of the fibronectin fibril networks is different whether platelets adhere to adhesion proteins from blood clots versus basement membranes

We finally asked whether our findings regarding Fn assembly could be generalized to specific thrombus sites. Besides Fn, we selected fibrin as the major clot component since spreading on fibrin but not fibrinogen supports Fn matrix assembly by platelets (Jaehyung Cho et al. 2005). Platelets on fibrin coatings showed the typical αIIbβ3-dependent contractile phenotype (Lickert et al. 2018) and formed Fn fibrils mainly localized in the periphery of the cell (Figure 6a). As on Fn coatings, Fn fibrils on fibrin were mainly pulled along the apical membrane (Figure 6c) and only a moderate increase in fibril length and thickness were noted (Supplementary Figure S8). Besides laminin, collagen type IV (Col4) is the most abundant component of the vascular basement membrane. Fn fibrils on Col4 coatings (Figure 6b) recapitulated the architecture on laminin with respect to their anchorage beneath the cell (Figure 6c), length and thickness (Supplementary Figure S8).

**Figure 6.**
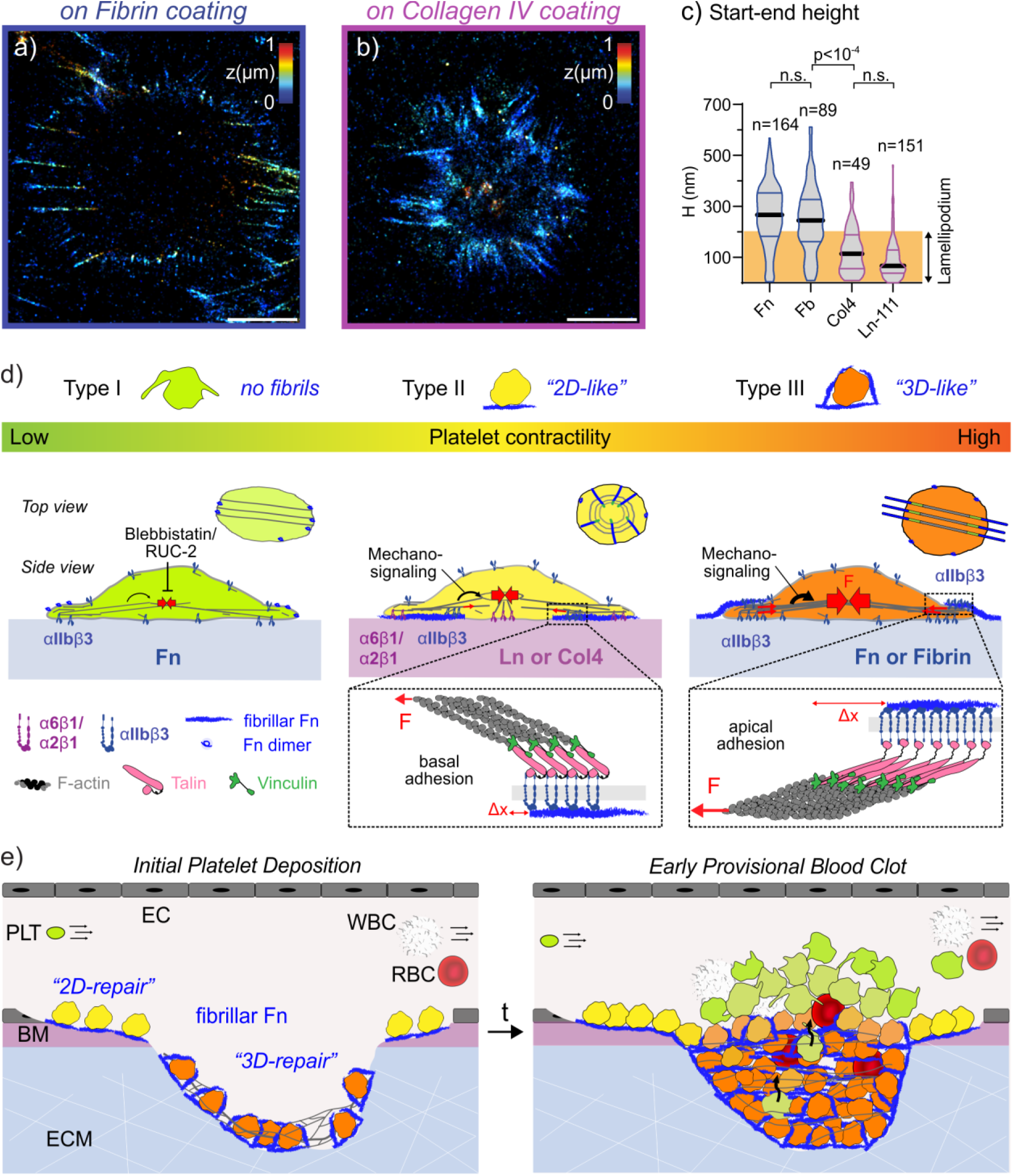
Fibronectin assembly of platelets on adhesion proteins of the provisional clot and the basement membrane. **(a)** 3D dSTORM image of pFn647-enriched Fn fibril assembly by a single platelet seeded on fibrin (Fb, binds αIIbβ3). See also Supplementary Movie S1. Scale bar 2 µm. **(b)** 3D dSTORM image of pFn647-enriched Fn fibril assembly by a single platelet seeded on collagen type IV (Col-4, binds α2β1). Scale bar 2 µm. **(c)** Systematic comparison of fibril start-end height between Fn (blue), Fb (blue), Col-4 (magenta) and Ln-111 (magenta). Data were pooled in addition to Figure 2 from two healthy human donors (31-33 years) and were compared with a non-parametric Kruskal-Wallis rank test with post-hoc Dunn test to make (multiple) comparisons. Integrin mechanosensing of ECM proteins triggers dimensionality of the first deposited ECM network. Platelets spread on Fn and Fb (right) generate high traction forces (red arrows and dark red-yellow) and Fn fibrils are aligned to polarized actin bundles (top view). Mechanosignaling (black arrow) through αIIbβ3 (blue) instructs Fn fibril anchorage along the apical membrane of platelets and the mechanomolecular strain induces a spatial offset between vinculin (green) and Fn (see inset). On Ln or Col4 (middle), platelets contract less strong (yellow) and form radial oriented Fn fibrils (top view) pulled beneath their basal side with a small spatial offset between vinculin and Fn (side view, inset). Dose-dependent inhibition of myosin IIa and the αIIbβ3 (left) further reduces platelet contractility (green) and prevents Fn fibril formation. **(e)** Based on these three phenotypes, we propose a model for the initial thrombus formation under flow. After rupture of the vessel wall, platelet adhesion to different ECM proteins tunes platelet contractility and orchestrates a distinct repair process. Platelet deposition on basement membrane (BM) proteins leads to a “2D-like” repair process with a less stretched planar Fn network. Platelets in contact with clot proteins (Fn and Fb) strongly contract and assemble a “3D-like” Fn network under high tension in the core of the clot and push less contractile platelets towards the thrombus surface. As a result, the fibrin network in the clot gets interlaced with Fn fibrils.

## DISCUSSION

How the local multifactorial biochemical and mechanical cues are sensed by platelets and direct their response to allow for a spatially well-organized repair of injured vessel walls versus scaring while maintaining mechanical integrity remains elusive. We discovered here by super-resolution microscopy that platelets can form fibrillar adhesions as fibroblasts do (Figures 1,5), but that they exploit αIIbβ3 rather than α5β1 integrins to drive Fn fibrillogenesis (Figures 3). For fibroblasts it is well established that mechanotransduction processes require the cooperation of mechano-sensing structural proteins, including talin, vinculin and tensin that connect the contractile cytoskeleton to the integrin tails and get stretched by forces, and of additional signalling molecules whose phosphorylation triggers downstream cell signalling programs, including FAK, Src and paxillin (Kechagia, Ivaska, and Roca-Cusachs 2019), yet it was not known that platelets can form fibrillar adhesions too. It is thus remarkable that platelets not only orchestrate thrombus formation and contraction, but also direct the initial tissue repair process by assembling Fn nanofibrils (Figures 1,2,6). This *de novo* ECM fibrillogenesis by platelets is in line with previous observations made at lower optical resolution on platelets alone (Olorundare et al. 2001; Jaehyung Cho et al. 2005; J. Cho and Mosher 2006a; Jaehyung Cho and Mosher 2006) or in blood clots (Burkhardt et al. 2016), and suggests that fibrillar Fn might play an overlooked role in the formation and remodeling of the hemostatic plug and the guidance of cell infiltration from the surrounding.

Our findings taken together with existing literature can be summarized in a new mechano-regulated thrombus formation model (Figure 6 d,e): we now propose the existence of at least three different platelet phenotypes, defined by either low, medium or high contractility. Each subtype serves contractile-specific functions, including their ability to sense their ECM environment and to respond by either being procoagulant or assembling *de novo* ECM. These three contractile phenotypes are not directly congruent with the well-known platelet phenotypes defined by α-/δ-granule secretion or PS exposure (van der Meijden and Heemskerk 2019). On one hand, more than one biochemical phenotype has low contractility, such as resting but also PS positive platelets (Zhang et al. 2018; Nechipurenko et al. 2019; Agbani et al. 2015). On the other hand, little is known about the contractility of platelets that have or have not undergone degranulation. The intricate feedback loops of both, platelet activation and platelet mechanosensing, and the cross-talk between them, demand a more comprehensive characterization of contractility in different platelet phenotypes to refine our proposed model.

Strongly contracting platelets push procoagulant or other non-contractile platelets to the thrombus surface (Nechipurenko et al. 2019) and thereby establish the basic core-shell architecture (Brass, Diamond, and Stalker 2016). Here we find that platelets in contact with proteins of the vessel wall are less contractile than when in contact with clot proteins (Figure 4). Platelet contractility drives the build-up of fibrillar adhesions (Figure 5) that coordinate the organization of distinct Fn fibril networks which either encage the platelets when adhering to Fn or fibrin (Figures 1,6) or are pulled beneath them on vascular basement membrane proteins (Figures 2,6). Extrapolated from our findings, the ECM microenvironment tunes platelet contractility at different thrombus locations (Figure 6e) suggesting that the platelets in contact with blood clot proteins play a far more active role in the wound site contraction. The fibrin- and Fn-rich extraluminal thrombus portion in penetrating injuries (Tomaiuolo et al. 2019; Wang et al. 2014) locally enhances compaction by maximizing platelet contraction, while platelets of medium contractility mediate the thrombus anchorage at the vascular basement membrane upon superficial injury. How additional chemical stimuli like gradients of platelet agonists or PS exposure could synergize with ECM proteins to up- (Olorundare et al. 2001) or downregulate (Zhang et al. 2018) platelet contractility and Fn fibrillogenesis needs further investigation. 2D spatial cues might guide invading cells to move along the plane that still contains basement membrane proteins or their fragments thus helping to restore the tissue barrier. The less stretched planar Fn fibril networks might further direct the 2D healing process by templating the reassembly of collagens (Kubow et al. 2015) for basement membrane repair, as during vascular morphogenesis (Hielscher et al. 2016) or in other tissues (Filla et al. 2017). An extensive Fn network at basement membranes is known to guide cell migration during development (Lu et al. 2020; Duband and Thiery 1982). We thus hypothesize that the platelet-deposited Fn fibrils at basement membranes could serve an analogous role during tissue repair. The progressive development of a tensed 3D network of Fn fibrils in the core might guide infiltrating cells or allow migrating platelets (Gaertner et al. 2017) to reposition themselves within a lesion. Our finding that the force which platelets apply per αIIbβ3 integrin to Fn fibrils equals the force which fibroblasts apply to the α5β1-Fn bonds, as probed here by the displacement of vinculin in fibrillar adhesion sites (Figure 5), is fascinating and likely of major physiological importance because it enables platelets to withstand the pulling forces of invading fibroblasts and hence ensures that the plug does not lose its mechanical stability during remodeling.

In platelets, αIIbβ3 and not α5β1 integrins are the sole drivers of Fn stretching and fibrillogenesis (Figure 3). Adhesion signaling of either α6β1 on laminin, α2β1 on Col4, or αIIbβ3 on Fn or fibrin is sufficient to drive αIIbβ3 to assemble Fn fibrils (Figures 1,2,6), in analogy to the pre-activation of α5β1 integrins by αvβ3 in fibroblasts (Zamir et al. 2000; Bharadwaj et al. 2017), the initiation of α5β1-containing sliding fibrillar adhesions by α3β1 or α2β1 integrins on Matrigel (Lu et al. 2020), and in accordance with previous observations (Olorundare et al. 2001; Jaehyung Cho et al. 2005; J. Cho and Mosher 2006b, 2006a; Jaehyung Cho and Mosher 2006). Adhesion signaling through αIIbβ3 is essential to develop full contractility (Figure 4), maximal stress fibers (Lickert et al. 2018), as well as a 3D apical anchorage of Fn fibrils as seen in fibroblasts (Zamir et al. 2000) (Figure 1) and a similar mechanomolecular strain within fibrillar adhesions (Figure 5). We for the first time derive a threshold of ∼1 nN per cell-substrate adhesion to activate αIIbβ3 integrins to form 3D fibrils (Figure 4). The ECM proteins from the vessel wall and the thrombus thus essentially instruct the platelets to adopt distinct phenotypes of medium and high contractility, respectively, probably serving different functions as guided by different levels of mechanosensitive integrin signaling (Strohmeyer et al. 2017; Chen et al. 2019). This mechanism in platelets is, probably due to their small size and the high abundance of αIIbβ3 integrins, different from fibroblasts. Even though α5β1 integrins compete with αIIbβ3 for their common ligand Fn and drive Fn fibrillogenesis in fibroblasts (Singh, Carraher, and Schwarzbauer 2010) (Figures 1,5), they surprisingly fail to rescue fibril assembly in platelets (Figure 3). The platelet kindlin 3 equally promotes integrin β1 and β3 clustering (Malinin et al. 2009) and talin binds either integrin, ruling out a deficiency for a specific adaptor protein for α5β1 in the myeloid lineage (Humphries et al. 2009). With ∼1’500 copies per cell (Zeiler, Moser, and Mann 2014) and a typical average force of ∼5 pN per integrin (Zhang et al. 2018), all α5β1 integrins together could only sustain 7.5 nN over prolonged time. This suggests that α5β1 integrins on platelets are merely outcompeted by the abundant αIIbβ3 during Fn fibrillogenesis. Although we did observe abundant tensin-1 immunostaining in platelets, in agreement with previous proteomic reports (Sun et al. 2007; Sabrkhany et al. 2018), tensin-1 did not localize to neither focal nor to fibrillar adhesion sites (Supplementary Figure S4). Since both, the NPLY motif of β3 integrins (Law, Nannizzi-Alaimo, and Phillips 1996) as well as myosin (Zimman et al. 2014; Solari et al. 2016), get phosphorylated upon thrombin-induced platelet aggregation, which might suggest a potential switch from talin to tensin in the contractile phenotype, further investigations are needed to delineate the functions of tensin-1 and of many other adaptor proteins in the fibrillar adhesions of platelets.

Beyond discovering that platelets can form fibrillar adhesions, our data reveal major new outside-in mechanisms by which the ECM identity of the microenvironment seen by platelets regulates their contractility, which vice versa adapts their phenotype leading to the built up of a heterogeneous thrombus and paving the way for cell invasion. In response to the ECM identity, which is rapidly changing in a fresh wound site, platelets change their contractility and the dimensionality of the first provisional ECM fibers which they assemble. This suggest new mechanisms by which the ECM microenvironment might direct location-specific responses, which upon integration, steer the early phases of a spatially well-organized tissue repair process. Beyond initiating blood coagulation and contracting blood clots, we now propose that the ability of platelets to interlace Fn fibers into the fibrin clot from early time points throughout its maturation might ensure that the mechanical stability of blood clots is maintained as the first provisional matrix gets degraded and replaced by newly formed tissue. We postulate that any factor that interferes with platelet Fn fibrillogenesis will thus contribute to bleeding disorders for example as resulting from myosin IIa mutations leading to MYH9 diseases (Nurden and Nurden 2008). Our findings have also direct relevance for how blood-biomaterial interactions (Burkhardt et al. 2016) instruct tissue repair processes and will probably guide the future development of wound healing scaffolds.

While our study investigated fundamental mechanisms of fibrillar adhesion formation by using isolated platelets, in order to specify the phenomenon, the integrins involved and how fibronectin fibrillogenesis is regulated by platelet contractility, we can only speculate here how the three platelet phenotypes might impact blood clot formation, contraction and the subsequent invasion of cells. Our mechanistic sketch (Figure 6e) though is in agreement with our previous observations of Fn fibrillogenesis and fibroblast invasion made on 2 and 24 hours old blood clots that had formed on titanium surfaces (Burkhardt et al. 2016).

## METHODS

### Reagents

Reagents were purchased from Sigma Aldrich, if not mentioned otherwise. Acid citrate dextrose (ACD) tubes (Sol. B, Vacutainer^®^, BD, Switzerland); coverslips (18 mm diameter, thickness 1.5; Hecht-Assistent, Germany); human fibrinogen (FG; F3879); human fibronectin (FN; purified from plasma as described previously (Früh et al. 2015)); human collagen type IV (Col4; C8374); murine laminin (LN-111; L2020); human laminin (LN-521, Biolaminin521 LN, BioLamina, Sweden); bovine serum albumin (BSA; 05470); Adenosine 5′-diphosphate sodium salt (ADP; A2754); thrombin from human plasma (T6884); RUC-2 (gift from Prof. B.S. Coller, New York University); Blebbistatin (B0560); mouse anti-integrin α5β1 (MAB1969); mouse anti-vinculin (V9131); unconjugated donkey anti-mouse or anti-rabbit IgG (Jackson Immunoresearch, USA); Alexa Fluor 647 NHS ester (A20006, ThermoFisher, USA); CF680 NHS ester (92139, Biotium, USA); Alexa Fluor 546 NHS ester (A20002, ThermoFisher, USA); DyLight 405 NHS ester (A46400, ThermoFisher, USA); goat anti-mouse CF680 (SAB4600361); Alexa Fluor 488 Phalloidin (A12379, ThermoFisher, USA); Alexa Fluor 647 Phalloidin (A22287, ThermoFisher, USA); human Factor XIII (Fibrogammin^®^ 1250, CSL Behring); Float-A-Lyzer G2 MWCO 20kD dialysis columns (Z726834-12EA); TI Prime (MicroChemicals, Germany); photoresist AZ 1505 (MicroChemicals, Germany); developer AZ 726 MIF (MicroChemicals, Germany); Sylgard 184 Silicone Elastomer (Dow Corning, USA); hard PDMS (PP2-RG07, Gelest, USA); Trichloro(1*H*,1*H*,2*H*,2*H*-perfluorooctyl)silane (448931), Pluronic^®^ F-127 (P2443).

### Labeling of plasma fibronectin (Fn) and BSA with fluorescent dyes

Purified Fn (in 1 M arginine in PBS) was stored at −80°C before use. The random labeling of surface accessible lysine residues of Fn was obtained by amid bond formation with fluorescent probes as described previously (Früh et al. 2015). In short, Fn was transferred into an amine labeling buffer (0.1 M NaHCO_3_ in PBS, pH 8.5) and incubated with 20-fold molar excess of Alexa Fluor 647 succinimidyl ester for 1 hour at room temperature. Free dye was removed and buffer exchanged to PBS. As measured by absorption, Fn-AF647 (denoted as pFn647) batch carried 10-15 dye molecules per molecule on average. For 2C STORM of pFn and F-actin, pFn was labeled with CF680 in an analogue way. For mPAD experiments, BSA was labeled with DyLight 405 and adhesion proteins were labeled with Alexa Fluor 488, resulting in 5-7 dyes per molecule.

### Platelets and sample preparation for dSTORM imaging

Fn, Ln and Col4 were coated with a concentration of 100 µg/ml (in PBS) over night at 4°C onto coverslips as described previously (Lickert et al. 2018). Cross-linked fibrin matrix was generated as described before (Jaehyung Cho et al. 2005) by mixing human fibrinogen (500 µg/mL); thrombin (3 unit/mL), FXIII (5 µg/mL), and CaCl_2_ (2 mM) in 0.5mL Tris-buffered saline (20 mM Tris-HCl, pH 7.4, and 150 mM NaCl and incubation of coverslips with this mixture overnight at 4 °C. All coverslips were washed thoroughly 3 times with PBS before use. Ethical approval was obtained from the Kantonale Ethikkommission Zurich (KEK-ZH-Nr. 2012- 0111) and RCSI Research Ethics Committee (REC1391 and REC1504) prior to the commencement of the study. All experiments were performed in accordance with relevant guidelines and regulations. Whole blood from healthy adult volunteers was collected in ACD tubes. Isolated washed platelets were resuspended in Tyrode’s buffer (TB) (TB containing 1.8 mM Ca^2+^,5 μM ADP, 90 µg/ml 10 µg/ml Fn-AF647, and, where appropriate, RUC-2, Blebbistatin or anti-integrin α5β1. After seeding on coverslips for 2 hours at 37 °C, platelets were rinsed with TB, fixed with 3% (w/v) paraformaldehyde (FA) in TB for 15 minutes and washed with PBS (see also Supplementary Information).

### Fabrication, optimization and imaging of the micropost substrates

Resist-coated silicon wafers were patterned with circles (Diameter: 1 µm; Center-to-center spacing: 2 µm) using 220 nm deep UV lithography (ABM, USA). Then, the resists were etched with a fluorine-based inductively coupled plasma (ICP) process (PlasmaPro100 Estrelas, Oxford Instruments, United Kingdom) to create 2.6 µm deep post structures. Master structures were replicated by creating elastomeric negative molds using a sandwich of spin-coated hard PDMS and Sylgard 184, and subsequent molding using hard PDMS on glass coverslips. The spring stiffness of posts was calculated as 34.51 nN/µm(Schoen et al. 2010). The top surface of the posts was then coated with adhesion protein (1:1 mixture of labeled/unlabeled) by contact printing after UV/Ozone activation. The remaining accessible mPAD surface was stained and passivated with fluorescent BSA (0.4 mg/mL) and then 0.5% (w/v) Pluronics F127. Washed platelets from healthy volunteers were seeded for 1 hr on the arrays and subsequently fixed with 3% (w/v) FA in PBS for 15 minutes. Samples were stained with phalloidin 647 and fluorescent z-stacks were acquired by confocal microscopy (Leica SP8, or Zeiss Examiner Z.1) at a pixel size of 60-70 nm. The deflection of posts was determined from the BSA channel. In short, positions of individual posts were determined by template matching and a radial symmetry fit. Positions of single posts were linked through slices of the z-stack. The lateral offset between slices was determined by a redundant cross-correlation of non-deflected posts in the region around cells and corrected. The deflection profile of posts was approximated by a c-spline, yielding deflection amplitude and directions of post tops. Forces per post were calculated by Hooke’s law using the spring stiffness. The median apparent force of posts in the region outside of cells was taken as the measurement resolution. The spreading area of cells was determined from outlines based on the F-actin stain, and the mean force per posts and the total force per cell were determined for each cell from the posts beneath it.

### dSTORM imaging of fibronectin and the cytoskeleton

During platelet seeding and spreading, the medium was supplemented with 10 µg/ml of pFn647 and 90 µg/ml unlabeled pFn that get incorporated into the Fn fibrils assembled by platelets, as previously observed for fibroblasts (Baneyx et al. 1999). Afterwards, samples were fixed and stained for the actin cytoskeleton by incubating with Alexa Fluor 488 phalloidin at 1:50 dilution for 1 h for epifluorescence. For 2C STORM of pFn and F-actin, Alexa Fluor 647 phalloidin was used instead. For 2C STORM of pFn and vinculin, samples were incubated overnight at 4°C with 1:60 dilution of anti-vinculin antibody in 3% (w/v) BSA. Next, samples were rinsed three times in PBS and incubated for 1 h with 1:60 dilution of anti-mouse CF680 in 3% BSA. After 3 washes in PBS, samples were post-fixed with 4% PFA in PBS for 15 min. A home-built set-up was used for single-molecule localization microscopy, as previously described (Früh et al. 2015). Fitting and analysis of dSTORM movies was performed using the software SMAP (developed by Dr. Jonas Ries, EMBL Heidelberg).

### Analysis of the fibril dimensions

Fn fibrils were analyzed as shown exemplary in Supplementary Figure S2. The dSTORM images were post-processed in SMAP and only z-localizations with a localization precision better than 100 nm further analyzed. Single fibers were manually marked from start to end by a line ROI which defined the x-coordinate. The length was taken as the length of the line. The diameter was defined as the full width half maximum (FWHM) of a Gaussian fit of the perpendicular line profile with 2 nm binning. The inclination and start-to end height were determined by a line fit to z-localizations binned into 20 intervals along x.

### Analysis of the lateral offset between vinculin and Fn fibrils at fibrillar adhesion sites

Lines were drawn along fibrillar adhesion sites from inside the cell towards to outside anchorage of the Fn fibrils. Fluorophore localizations in both channels (vinculin and Fn) that were within a 150 nm wide region around these lines were rotated to align them in the x direction. The start of the vinculin adhesion and of the Fn fibril were set to the 0.05 quantile of x-positions of localizations in the respective channel. This procedure allowed for a robust determination of the signal boundaries in the presence of background localizations (Supplementary Figure S9).

### Manual count of Fn fibrils

Fraction of platelets which assembled Fn were measured manually. At least thirty different field of views (48.1 × 48.1 µm^2^) were captured in the pFn channel on the epifluorescence microscope and platelets which assemble pFn fibrils (fibril length > 1 µm) and platelets which only deposit pFn (fibril length < 1 µm) were separately counted and the ratio determined.

## Acknowledgements

We thank Prof. Barry S. Coller for generously providing the RUC compounds and Dr. Jonas Ries (EMBL Heidelberg) for kindly providing software for the visualization of dSTORM data. We would like to acknowledge Lukas Braun for the helpful discussion during the manuscript preparation. This work was supported by ETH Zurich, by Swiss TransMed “Life Matrix” 33/2013 (V.V.), Wyss Zurich (V.V.), the Velux Stiftung (K.S.), as well as by the RCSI (I.S.), and has received funding from the European Union’s Horizon 2020 research and innovation programme under grant agreement No 747586 (I.S.).

## Author contributions

S.L., I.S. and V.V. laid out the major concepts of the study; M.A.B. performed initial fibrillogenesis experiments with platelets; S.L. and I.S. planned experiments; S.L., K.S. and M.K. performed all platelet spreading experiments; S.M.F. performed cell culture experiments with fibroblasts; S.L. with help of K.S. and J.L.M. performed super-resolution imaging for platelets, S.M.F. performed super-resolution imaging of fibroblasts; S.L. and I.S. wrote the script for fibril analysis and analyzed all super-resolution data; S.L fabricated the micropost arrays; S.L., M.K. and I.S. optimized the micropost arrays; S.L. and M.K performed, imaged and analyzed platelet traction force experiments; I.S. wrote the script for the traction forces analysis; S.L., I.S. and V.V. wrote the manuscript. All authors critically revised the report.

