## Supporting Information for "Platelets exploit fibrillar adhesions to assemble fibronectin matrix revealing new force-regulated thrombus remodeling mechanisms"

### 1 Supplementary Figures and Movies

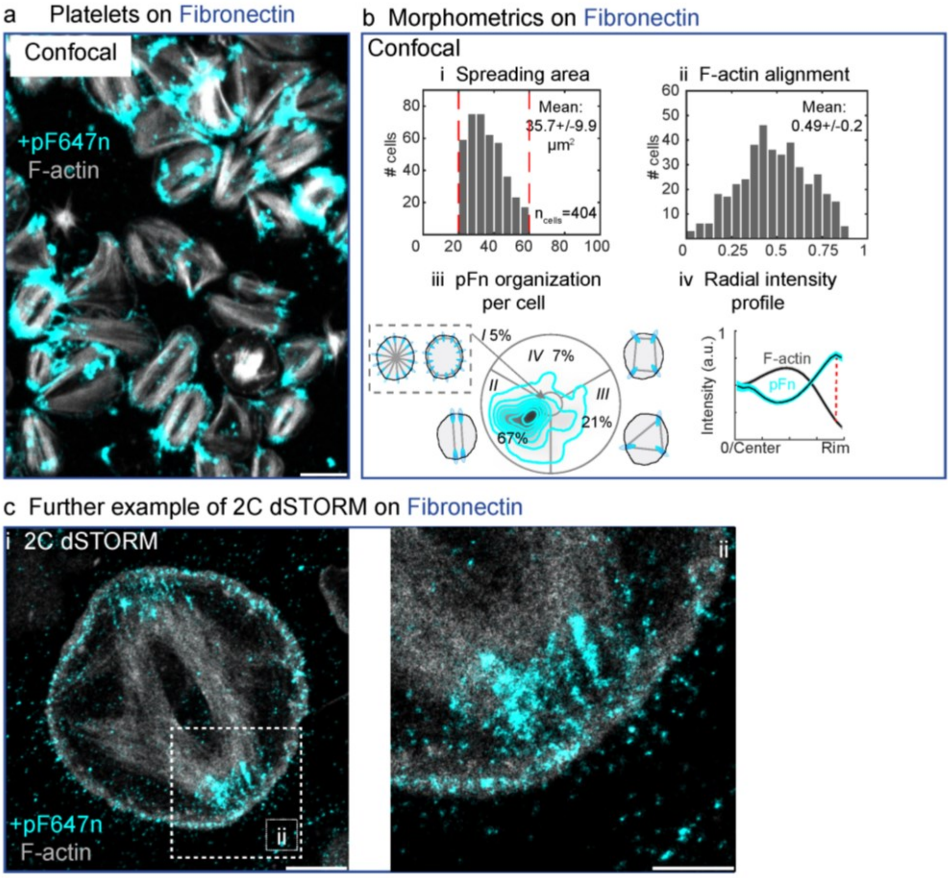

**Figure S1. Morphometry of platelet assembled fibronectin fibrils and 2C dSTORM of F-actin and pFn.**

(a) Cells are seeded on Fibronectin (Fn) in the presence of fluorescently labeled plasma Fibronectin (pFn647; cyan), fixed after 60 minutes, stained for F-actin (grey) and imaged using Confocal Laser Scanning Microscopy (CLSM). Scale bar 5 µm. (b) Morphometric analysis of platelets from one healthy donor with regards to (i) spreading area, (ii) alignment of F-actin, (iii) the distribution of pFn647 per cell, and (iv) the radial intensity for F-actin and pFn per cell. (c) Example of a dual color (2C) dSTORM image of a single platelet spread on Fn with supplemented pFn647 (cyan) during seeding and stained in addition for F-actin (gray). Scale bar 2 µm. (ii) Magnification of the boxed region in (i). Scale bar 1 µm.

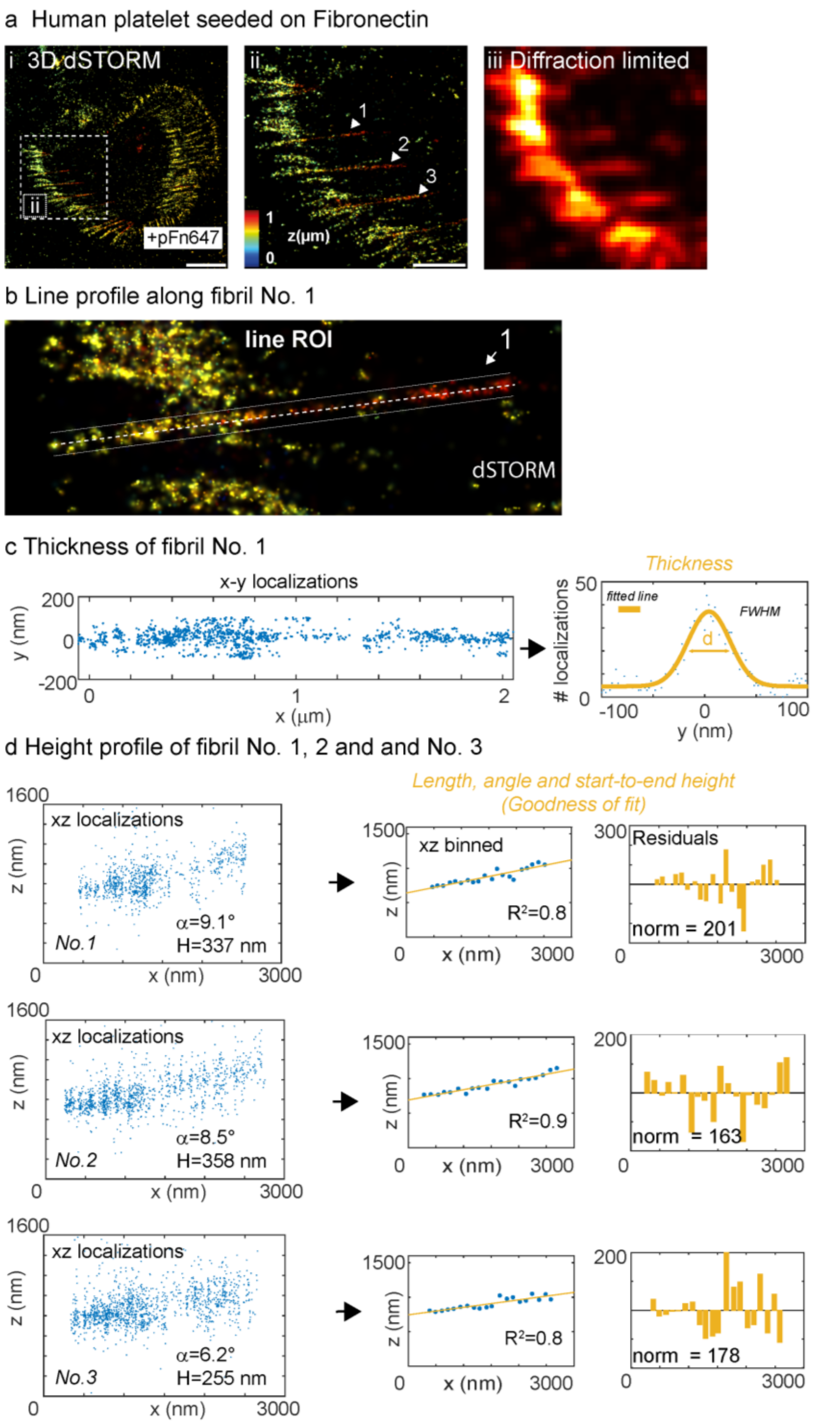

**Figure S2. Analysis of the spatial dimensions of Fn fibrils assembled by platelets seeded on fibronectin.**

**(a)** Fn fibril assembly by a single platelet spread on Fn with the addition of pFn followed by fixation and dSTORM imaging. The z-position is color-coded from blue (basal) to red (apical). Scale bar 2 µm. (ii) Magnification of the boxed region in (i) and each Fn fibril is marked arrow head. Scale bar 1 µm. (iii) Diffraction limited representation of (ii). **(b)** Manually dawn line ROI along fibril 1 from (a). **(c)** Determination of the fibril thickness of fibril 1 in (b) by binning the xy-localization (left, blue dots) within the line ROI perpendicular to the fibril direction and fitting by a Gaussian (right, yellow line) to derive the full width half maximum (FWHM) which serves as a measure for the fibril diameter/thickness. **(d)** Determination of the fibril start-to-end height for all marked fibrils in (a). The x-z localizations were extracted from the line ROI (left, blue dots), binned and fitted with a straight line (middle, yellow line). The goodness of the fit is checked by the coefficient of determination $R^2$ and by plotting the residuals (right, yellow bars).

a Basic demographics of healthy donors used in this study

| Donor | Age | Gender | Main figure |
|---|---|---|---|
| 1 | 33 | m | 1,2,4,6 |
| 2 | 31 | m | 1,2,3,5,6 |
| 3 | 32 | f | 1,2,6 |
| 4 | 33 | m | 2,3,5 |
| 5 | 35 | f | 1,2,6 |
| 6 | 27 | f | 2,5 |
| 7 | 31 | m | 1,2,3,4 |
| 8 | 33 | m | 4 |

b Donor-to-Donor variation for platelets seeded on Fibronectin

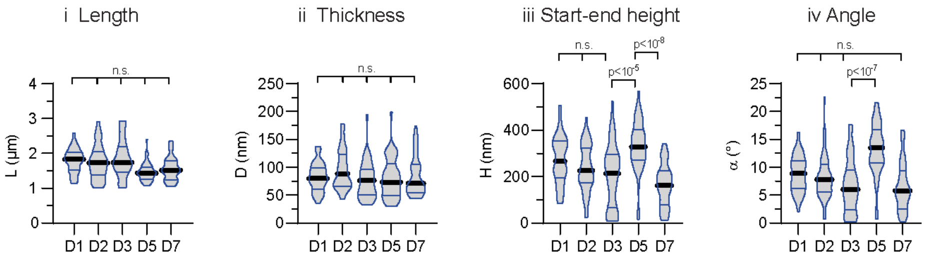

c Day-to-day variation (D2) for the same donor for platelets seeded on Fibronectin

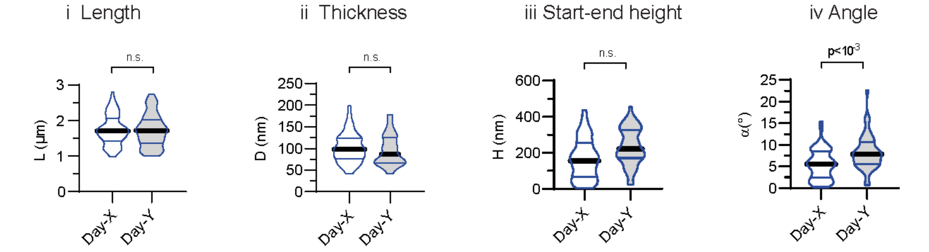

**Figure S3. Donor-to-donor and day-to-day variations of Fn fibril dimensions.**
**(a)** Platelets were obtained for this study from healthy donors between 25-35 years of age. **(b)** Comparison of Fn fibril dimensions from five different donors with respect to (i) length L, (ii) thickness D, (iii) start-end height H, and (iv) angle $\alpha$. Only fibrils longer than 1 µm were analyzed. The distribution of the data are depicted using violin plots showing the median (thick black line) and interquartile ranges (thin colored lines). Multiple comparisons are made using a non-parametric Kruskal-Wallis rank test with post-hoc Dunn test. **(c)** Comparison of Fn fibril dimensions using blood from the same donor but different withdrawals more than 2 weeks apart. Data were compared with an unpaired two-tailed Mann-Whitney test. Adjusted p-values below 0.001 were accepted as highly significant (Annotation: n.s. – not significant).

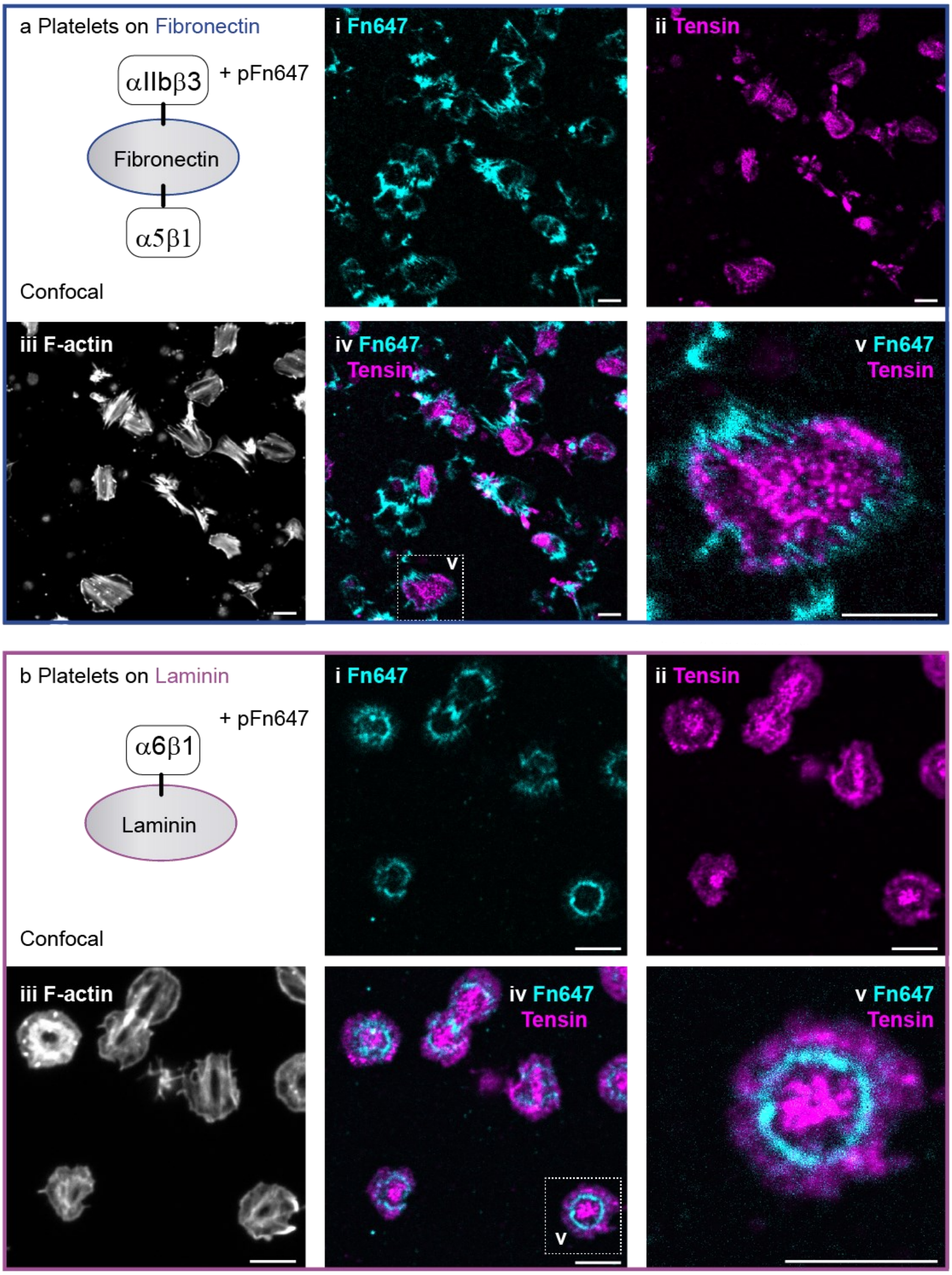

**Figure S4. Immunostaining of tensin-1 in platelets seeded on fibronectin and laminin coatings.**
**(a,** blue box**)** Cells are seeded on Fibronectin (Fn) in the presence of fluorescently labeled plasma Fibronectin (pFn647; cyan, i), fixed after 60 minutes, stained for tensin-1 (magenta, ii), F-actin (grey, iii). pFn signal and the tensin stain are overlaid (iv) and the boxed region is magnified (v). **(b,** magenta box**)** Cells are seeded on Laminin-111 (Ln) in the presence of fluorescently labeled plasma Fibronectin (pFn647; cyan, i), fixed after 60 minutes, stained for tensin-1 (magenta, ii), F-actin (grey, iii). pFn signal and the tensin stain are overlaid (iv) and the boxed region is magnified (v). All images are obtained by using Confocal Laser Scanning Microscopy (CLSM). Scale bar 5 µm.

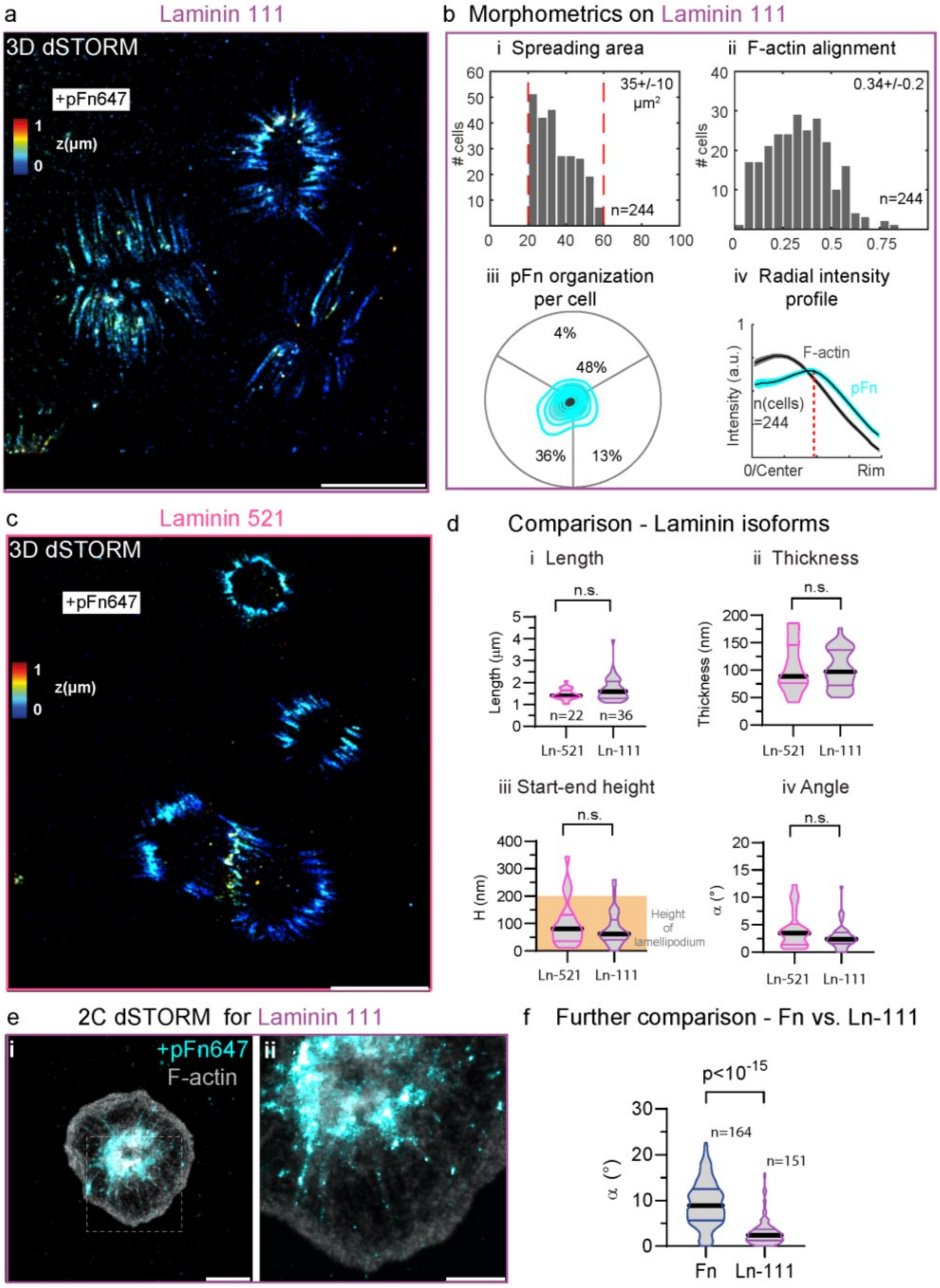

**Figure S5. Fn fibril assembly by platelets seeded on different laminin isoforms**.

**(a)** Fn fibril assembly by a single platelet spread on laminin-111 (Ln-111) with the addition of pFn followed by fixation and dSTORM imaging. The z-position is color-coded from blue (basal) to red (apical). Scale bar 5 µm. **(b)** Morphometric analysis of platelets from one healthy donor with regards to (i) spreading area, (ii) alignment of F-actin, (iii) the distribution of pFn647 per cell, and (iv) the radial intensity for F-actin and pFn per cell. **(c)** Fn fibril assembly by a single platelet spread on laminin-521 (Ln-521) with the addition of pFn followed by fixation and dSTORM imaging. Scale bar 5 µm. **(d)** Comparison of Fn fibril dimensions from platelets seeded on the different laminin isoforms. Data were compared with an unpaired two-tailed Mann-Whitney test. **(e)** Further example of a dual color (2C) dSTORM image of a single platelet seeded on Ln-111 with supplemented pFn647 (cyan) and stained in addition for F-actin (gray) after fixation. Scale bar 2 µm. (ii) Magnification of the boxed region in (i). Scale bar 1 µm. **(f)** Comparison of the tilt towards the substrate of the Fn fibrils from platelets spread on Ln-111 and Fn. Data were compared with an unpaired two-tailed Mann-Whitney test.

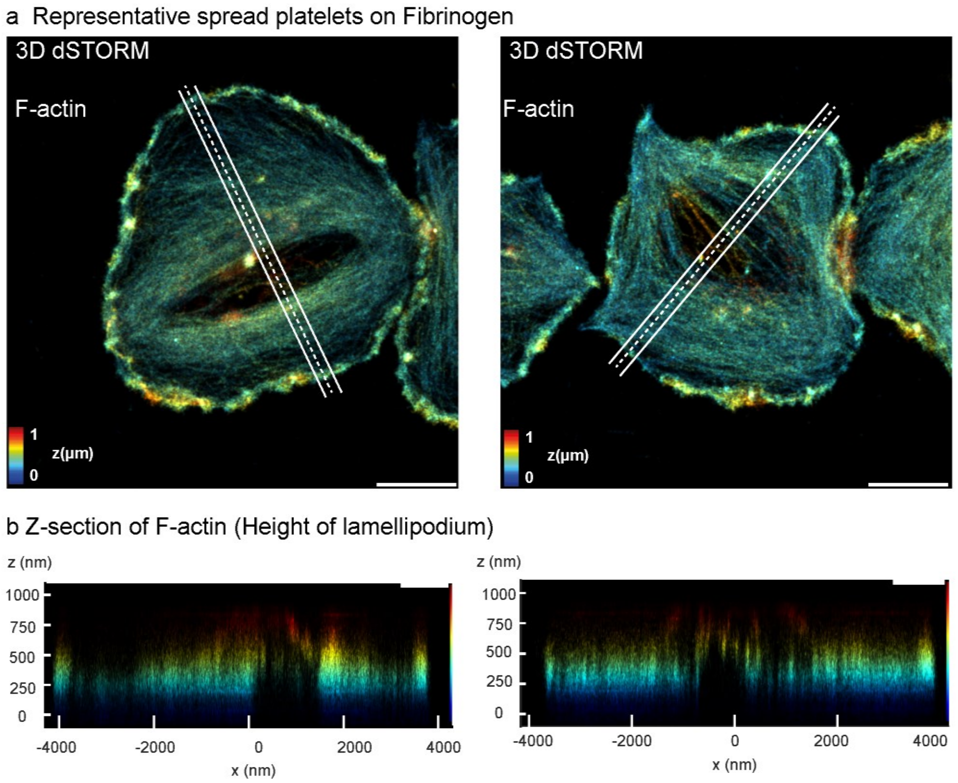

**Figure S6. Height of the lamellipodium of spread platelets.**
**(a)** Two representative dSTORM images of platelets spread on fibrinogen (Fg) and stained for F-actin. The z-position is color-coded from blue (basal) to red (apical). Scale bar 2 µm. **(b)** The line profile in (a) over the whole cell is depicted in a side view representation for both cells and the region at the cell edge represents the height of the lamellipodium. The height was estimated to be ~200 nm.

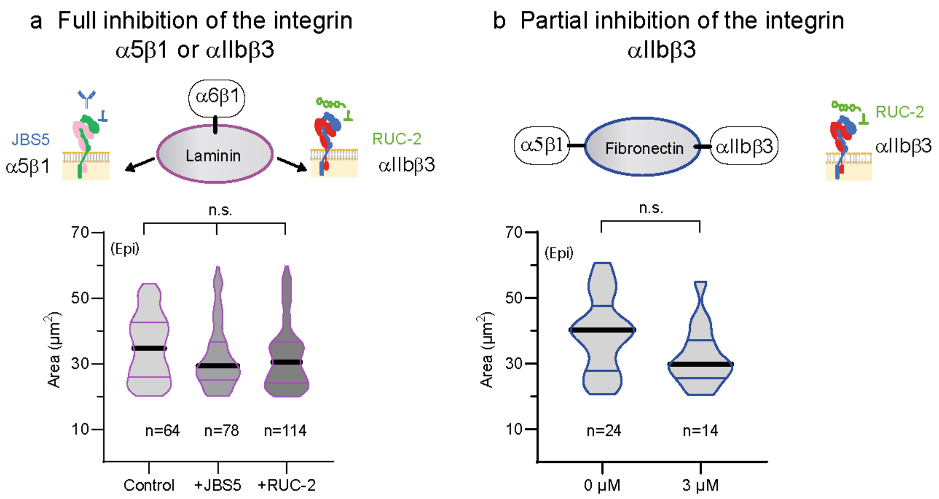

**Figure S7. Spreading area of platelets in the presence of different inhibitors.**
**(a)** Comparison of the spreading area of platelets seeded on Ln-111 in the presence of saturating concentrations of inhibitors for integrin α5β1 (20 mg/mL JBS5) or αIIbβ3 (100 μM RUC-2). Spreading area was determined from epifluorescence F-actin images. Multiple comparisons are made between data using a non-parametric Kruskal-Wallis rank test with post-hoc Dunn test. **(b)** Comparison of the spreading area of platelets on Fn partially blocked for αIIbβ3 (3 μM RUC-2). Data were compared with an unpaired two-tailed Mann-Whitney test.

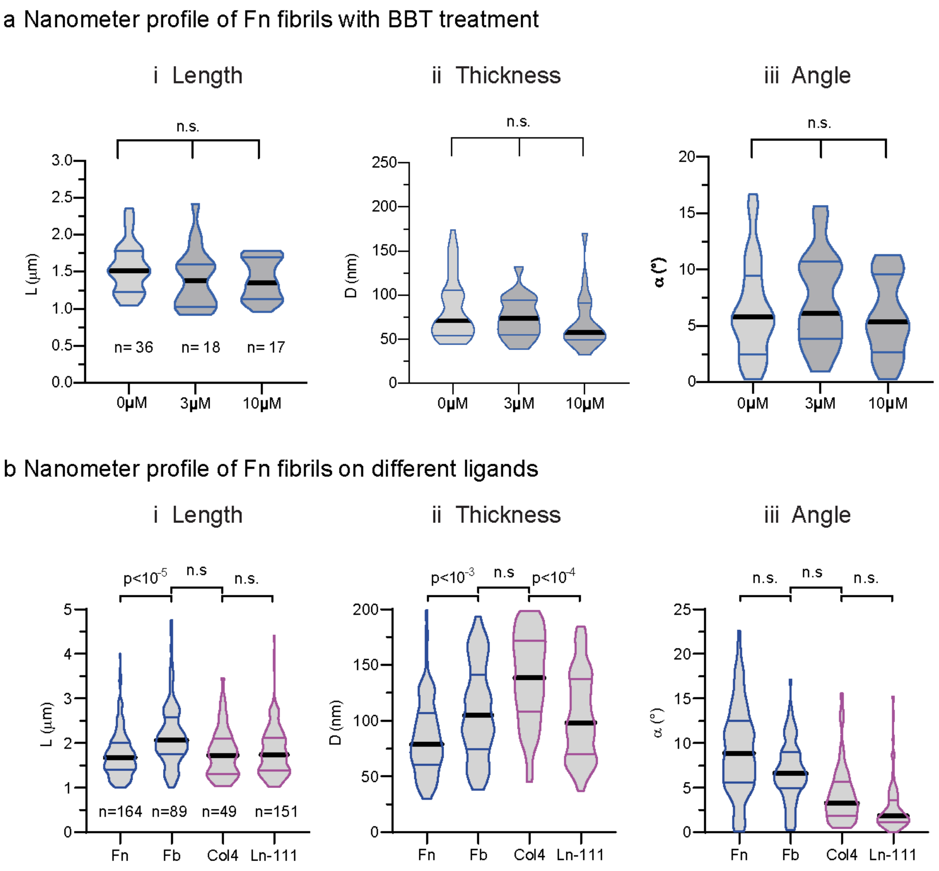

**Figure S8. Dimensions of Fn fibrils assembled by platelets treated with blebbistatin or spread on different ligands.**

**(a)** Comparison of (i) length L, (ii) thickness D, or (iii) angle $\alpha$ of Fn fibrils from platelets spread on Fn and treated dose-dependently with blebbistatin. **(b)** Comparison of Fn fibril dimensions from platelets spread on different ligands, including fibronectin (Fn), fibrin (Fb), collagen type IV (Col4), or laminin (Ln-111). Data were compared with a non-parametric Kruskal-Wallis rank test with post Dunn test to make (multiple) comparisons.

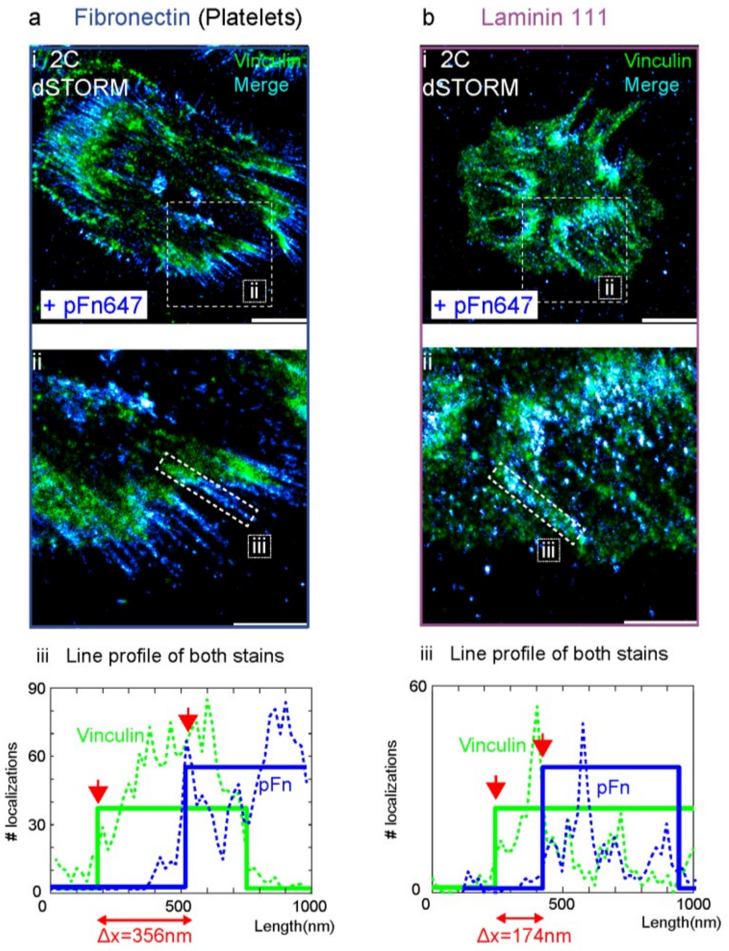

**Figure S9. Experimental determination of the spatial offset between vinculin and pFn.**
**(a)** A representative platelet spread on Fn with supplemented pFn (blue) and stained in addition for vinculin (green). Scale bar 2 µm. (ii) Magnification of the boxed region of Fn fibrils at the cell edge in (i). (iii) The start of the vinculin patch (green) or Fn fibril (blue) was determined from the 0.05 quantile of localizations along manually drawn line profiles. Shown is one example for the boxed region in (ii). **(b)** A representative platelet spread on Ln-111 with supplemented pFn during seeding and stained in addition for vinculin. Scale bar 2 µm. (ii) Magnification of the boxed region of Fn fibrils at the cell edge in (i). (iii) The start of the vinculin patch (green) or Fn fibril (blue) was determined from the 0.05 quantile of localizations along manually drawn line profiles. Shown is one example for the boxed region in (ii).

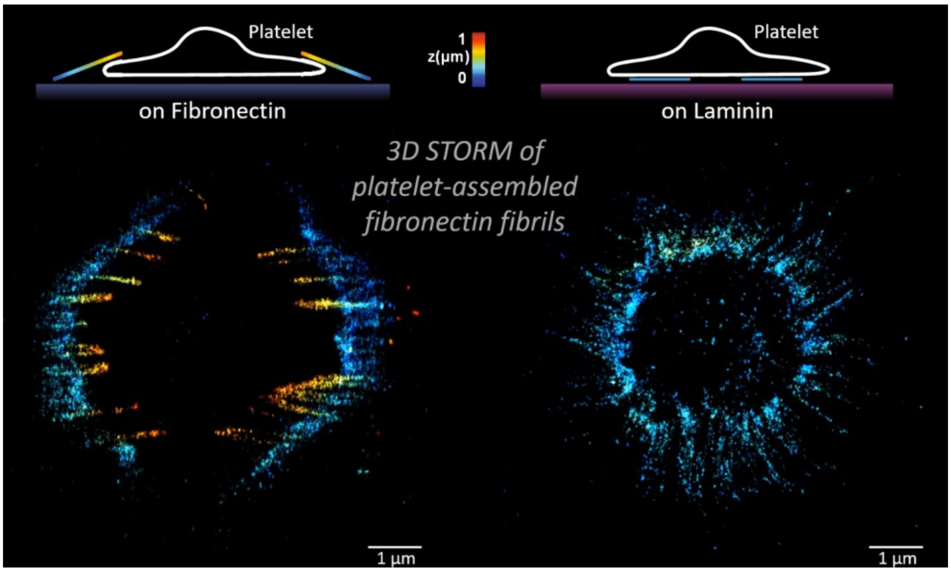

**Movie S1. Animation of 3D dSTORM images of platelet-assembled fibrinectin fibrils on fibronectin versus laminin coatings.**

3D dSTORM comparison of pFn647-containing fibronectin fibrils assembled by platelets on fibronectin coatings (left) versus on Laminin-111 coatings (right). Data are the same as in Figure 2 a+d. Schematics (top) are not to scale.

### 2 Additional experimental methods

#### 2.1 Platelet isolation

Platelets were isolated not later than 4 hours after blood withdrawal. Whole blood was centrifuged in the collection tubes at 180 g for 15 min at room temperature. 1.5 mL of platelet rich plasma (PRP) was collected into 2 mL tubes, 400 µL of ACD solution (dextrose 1.47% (w/v); tri-sodium citrate dihydrate 1.32% (w/v); anhydrous citric acid 0.48% (w/v)) was added, contents were gently mixed by inversion, and centrifuged at 900 g for 5 min at room temperature. Supernatant was removed and platelet pellet was re-suspended in 500 µL of pre-warmed (at 37°C) Tyrode's buffer (TB: 134 mM NaCl, 12 mM $NaHCO_3$, 2.9 mM KCl, 0.34 mM $Na_2HPO_4$, 1 mM $MgCl_2$, 10 mM HEPES, pH 7.4). 20 µL of re-suspended platelet pellet was added to each well in 700 µL seeding buffer (Tyrode's buffer containing 1.8 mM $CaCl_2$, 5 µM ADP, 10 µg/mL Fn-AF647, 90 µg/mL unlabeled Fn). Seeded platelets were incubated in the dark at 37 °C for 2 hours. Next, samples were washed 3 times with TB and fixed with 3% paraformaldehyde in TB for 15 min. Samples were washed three times with PBS and stored at 4°C.

For traction force measurements, the isolation protocol was slightly modified by adding prostaglandin E (PGE)-1 to the PRP at final concentration 1 µM, by including 0.05 U/mL apyrase in the wash buffer, and repeating the resuspension step another time. Platelet were counted on a hematometer (Sysmex). 8 million platelets were added to one 12-well chamber that contained the mPADs on a 20 mm coverslip and 800 µL seeding buffer (TB containing 1.8 mM $CaCl_2$, 5 µM ADP), gently mixed, and incubated at 37°C for 1 hour.

#### 2.2 Morphometrics of plasma Fibronectin distribution (pFn) and confocal imaging

Coverlips were mounted in a chamber (Chamlide; Live Cell Instruments, South Korea) on a confocal laser scanning microscope (SP8; Leica Microsystems, Germany) using a 63x oil immersion objective and 2x zoom (resulting in a field of view of 123 x 123 µm at 60.1 nm pixel size) and excitation at 488 nm and 647 nm. For statistical analysis of platelet morphology and the pFn distribution, between 200-300 cells were recorded per condition at 6-8 different field of views on the sample.

Morphometric image analysis was performed as previously described[1]. In short, outlines and morphological features of single platelets were automatically extracted from fluorescence images by different morphological and filtering operations. The outline yielded the single cell spreading area. The F-actin alignment was quantified by the average of the cosine of the local orientation of actin/Fn fibrils relative to their mean orientation. The radial distribution profile of the pFn staining was subjected to a Fourier decomposition and the resultant amplitudes were used to assign a characteristic morphology (isotropic, bipolar, triangular) to the cell. Then, the distribution of many platelet morphologies was depicted as a contour plot.
